# A confirmation bias in perceptual decision-making due to hierarchical approximate inference

**DOI:** 10.1101/440321

**Authors:** Richard D. Lange, Ankani Chattoraj, Jeffrey M. Beck, Jacob L. Yates, Ralf M. Haefner

**Affiliations:** Brain and Cognitive Sciences, University of Rochester, Rochester, NY 14627, USA.; Computer Science, University of Rochester, Rochester, NY 14627, USA.; Department of Neurobiology, Duke University, Durham, NC 27708, USA.; Department of Neurobiology, University of Pennsylvania, Philadelphia, PA 19104; Department of Biology, University of Maryland, College Park, MD 20742

## Abstract

Making good decisions requires updating beliefs according to new evidence. This is a dynamical process that is prone to biases: in some cases, beliefs become entrenched and resistant to new evidence (leading to primacy effects), while in other cases, beliefs fade over time and rely primarily on later evidence (leading to recency effects). How and why either type of bias dominates in a given context is an important open question. Here, we study this question in classic perceptual decision-making tasks, where, puzzlingly, previous empirical studies differ in the kinds of biases they observe, ranging from primacy to recency, despite seemingly equivalent tasks. We present a new model, based on hierarchical approximate inference and derived from normative principles, that not only explains both primacy and recency effects in existing studies, but also predicts how the type of bias should depend on the statistics of stimuli in a given task. We verify this prediction in a novel visual discrimination task with human observers, finding that each observer’s temporal bias changed as the result of changing the key stimulus statistics identified by our model. By fitting an extended drift-diffusion model to our data we rule out an alternative explanation for primacy effects due to bounded integration. Taken together, our results resolve a major discrepancy among existing perceptual decision-making studies, and suggest that a key source of bias in human decision-making is approximate hierarchical inference.

## Introduction

Human decisions are known to be systematically biased, from high-level planning and reasoning to low-level perceptual decisions [65, 42]. Decisions are especially difficult when they require synthesizing multiple pieces of noisy or ambiguous evidence for or against multiple alternatives [13, 6, 26, 18]. Perceptual decision-making studies across multiple species and sensory modalities have exposed systematic biases that differ in ways that are not well understood. Here, we focus on temporal biases, which range from over-weighting early evidence (a primacy effect) to over-weighting late evidence (a recency effect) (Figure 1a) even in situations when an equal weighting of evidence would be optimal. Despite seemingly comparable tasks, existing studies are surprisingly heterogeneous in the biases they find: some report primacy effects [33, 43, 72], some find that information is weighted equally over time [76, 11, 54], and some find recency effects [19] without a clear pattern emerging from the data.

**Figure 1:**
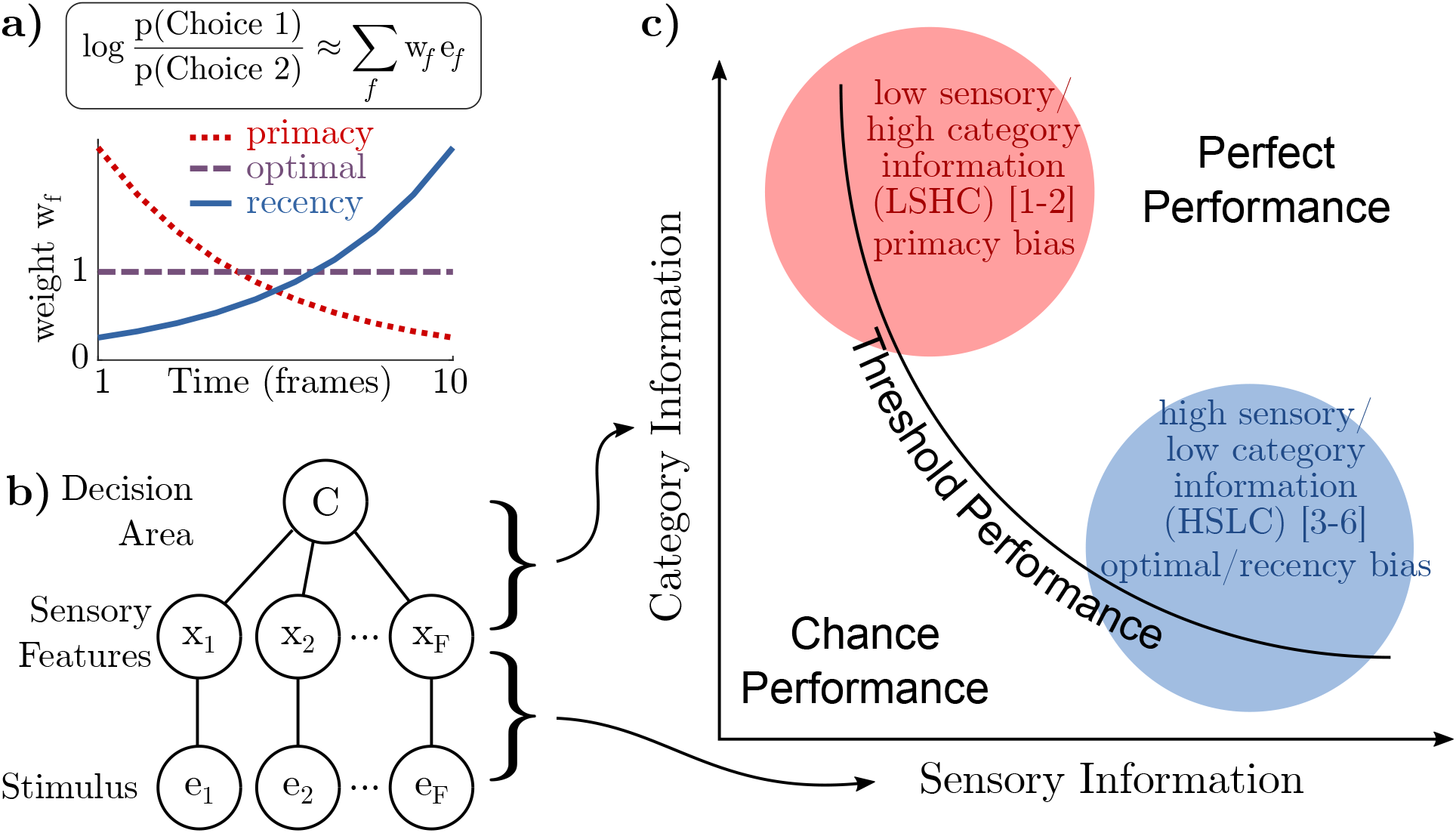
Differences in “sensory information” and “category information” can explain differences in temporal biases reported by earlier studies. **a)** A observer’s “temporal weighting strategy” is an estimate of how their choice is based on a weighted sum of each frame of evidence *e_f_* (more precisely, a weighted sum of the log odds at each frame). Three commonly observed motifs are decreasing weights (primacy), constant weights (optimal), or increasing weights (recency). **b)** Information in the stimulus about the category may be decomposed into information in each frame about a sensory variable (“sensory information”) and information about the category given the sensory variable (“category information”). **c)** Category information and sensory information may be manipulated independently, creating a two-dimensional space of possible tasks. Any level of task performance can be the result of different combinations of sensory and category information. A qualitative placement of previous work into this space separates those that find primacy effects in the upper-left (low sensory/high category information or LSHC regime) from those that find recency effects or optimal weights in the lower right (high sensory/low category information or HSLC regime). Numbered references are: [1] Kiani et al. 2008, [2] Nienborg and Cumming 2009, [3]	Brunton et al. 2013, [4] Wyart et al. 2012, [5] Raposo et al. 2014, [6] Drugowitsch et al. 2016.

Existing models propose mechanisms for either primacy [33] or recency [25] effects alone, or are flexible enough to account for either type of bias [13, 6, 75, 64, 11, 19], but none identifies or predicts the factors that cause one bias or the other to appear in a given context. All of these models are based on a variant of the classic drift-diffusion model [26]. For example, Kiani et al. proposed that evidence integration stops when an internal bound is reached, even during fixed stimulus duration tasks. Averaged over many trials in which the bound is reached at different times, this leads to a primacy effect. Alternatively, a primacy effect is also expected if evidence integration is implemented by an attractor network [74, 75, 64]. However, neither of these mechanisms can account for recency effects. On the other hand, including a “forgetting” term in the updating of the decision-variable leads to a recency effect [13, 6, 64, 25]. The analysis by Glaze et al. shows that a recency bias is optimal in a volatile environment, but such mechanisms cannot explain primacy effects. Deneve’s normative analysis predicts that primacy and recency should depend on trial-by-trial changes in difficulty, while Prat-Ortega et al. find that primacy and recency can change as a function of the variability of the input to a attractor-based decision-circuit. However, neither account alone, or in combination, can explain the differences found across experiments. It is thus an open question whether the disparate biases observed empirically are due to differences in species, sensory modalities, training, experimental design, or individual observers.

Here, we propose a new model that not only accounts for the existing findings in the literature, but also predicts which key aspect of the stimulus determines the specific temporal bias shown by an observer. Our model extends classic ideal observer models to the hierarchical case by explicitly including the intermediate sensory representation. This reveals that task difficulty is modulated by two distinct types of information: the information between the stimulus and sensory representation (“sensory information”), and the information between sensory representation and category (“category information”) (Figure 1b-c). We show that approximate inference in such a model predicts characteristic temporal biases in a way that can explain prior empirical findings. Furthermore, our model makes a critical prediction: that the temporal bias of an individual observer should change from primacy to recency as the balance in the types of information is changed. We verify this critical prediction of our model using newly collected data from a novel pair of visual discrimination tasks. Finally, we perform a quantitative model comparison demonstrating that inference dynamics, not a finite integration bound, explain our observers’ biases, consistent with our theory.

## Results

### “Sensory Information” vs “Category Information”

Normative models of decision-making in the brain are typically based on the idea of an ideal observer, who uses Bayes’ rule to infer the most likely category on each trial given the stimulus. On each trial in a typical task, the stimulus consists of multiple “frames” (by “frames” we refer to independent pieces of evidence that are not necessarily visual). If the stimulus or evidence in each frame, *e_f_*, is independent, then information about the category *C* can be combined by the well-known process of summing the log odds implied by each piece of evidence [68, 6, 26]:

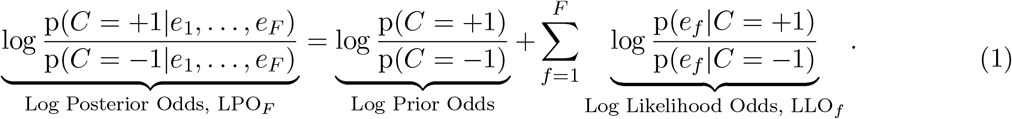

The ideal observer updates their current belief about the correct category by adding the information provided by the current evidence: LPO_*f*_ = LPO_*f*-1_ + LLO_*f*_.

In the brain, however, a decision-making area cannot base its decision on the externally presented stimulus, *e_f_*, directly, but must rely on intermediate sensory features, which we call *x_f_* (Figure 1b). Accounting for the intervening sensory representation implies that LLO_*f*_ cannot be computed directly, but only in stages. The information between the stimulus and category (*e_f_* to *C*) is therefore partitioned into two stages: the information between the stimulus and the sensory features (*e_f_* to *x_f_*), and the information between sensory features and category (*x_f_* to *C*). We call these “sensory information” and “category information,” respectively (Figure 1b). These two kinds of information define a two-dimensional space in which a given task is located as a single point (Figure 1c). For example, in a visual task, each *e_f_* would be the image on the screen while *x_f_* could be the instantaneous orientation or motion direction.

An evidence integration task may be challenging either because each frame is perceptually unclear (low “sensory information”), or because the relationship between sensory features and category is ambiguous in each frame (low “category information”). Consider the classic dot motion task [41] and the Poisson clicks task [11], which occupy opposite locations in the space. In the classic low-coherence dot motion task, observers view a cloud of moving dots, a small percentage of which move “coherently” in one direction. Here, sensory information is low since the percept of net motion is weak on each frame. Category information, on the other hand, is high, since knowing the true net motion on a single frame would be highly predictive of the correct choice (and of motion on subsequent frames). In the Poisson clicks task, on the other hand, observers hear a random sequence of clicks in each ear and must report the side with the higher rate. Here, sensory information is high since each click is well above sensory thresholds. Category information, however, is low, since knowing the side on which a single click was presented provides only little information about the correct choice for the trial as a whole (and the side of the other clicks). When frames are sequential, another way to think about category information is as “temporal coherence” of the stimulus: the more each frame of evidence is predictive of the correct choice, the more the frames must be predictive of each other, whether a frame consists of visual dots or of auditory clicks. Note that our distinction between sensory and category information is different from the well-studied distinction between internal and external noise; in general, both internal and external noise will reduce the amount of sensory and category information.

In general, sensory and category information depends on the nature of the sensory features x relative to *e* and *C*, and those relationships depend on the sensory system under consideration. For instance, a high spatial frequency grating may contain high sensory information to a primate, but low sensory information to a species with lower acuity. Similarly, when “frames” are presented quickly, they may be temporally integrated, with the effect of both reducing sensory information and increasing category information.

Qualitatively placing prior studies in the space spanned by these two kinds of information results in two clusters: the studies that report primacy effects are located in the upper left quadrant (low-sensory/high-category or LSHC) and studies with flat weighting or recency effects are in the lower right quadrant (high-sensory/low-category or HSLC) (Figure 1c, see Supplemental Text for justifications of placements). This provides initial empirical evidence that the trade-off between sensory information and category information may underlie differences in temporal weighting seen in previous studies. Unfortunately, since our placement of prior studies is only qualitative this observation only constitutes weak evidence in favor of this hypothesis. However, this hypothesis makes the strong prediction that a simple change in the stimulus statistics corresponding to sensory and category information, while holding everything else constant, should change the temporal weighting found in these previous studies (predictions provided in Supplemental Table S1). Below we will present new data from an experiment in which we did exactly that and found that biases indeed shifted from primacy to optimal/recency as predicted.

### Approximate hierarchical inference explains transition from primacy to recency

If stimuli were processed by the brain in a purely feedforward fashion, then a decision-making area could simply integrate the evidence in sensory features (*x_f_*) directly. This is consistent with some theories of inference in the brain in which sensory areas represent a likelihood function over stimuli [38, 3, 51, 69]. However, activity in sensory areas does not rigidly track the stimulus, but is known to be influenced by past stimuli [77, 37], as well as by feedback from the rest of the brain [24, 32]. In fact, the intermediate sensory representation is itself often assumed to be the result of an inference process over latent variables in an internal model of the world [39, 36, 78]. This process is naturally formalized as *hierarchical inference* (Figure 2a) in which feedforward connections communicate the likelihood and feedback communicates the prior or other contextual expectations, and sensory areas combine these to represent a posterior distribution [20, 51, 23, 61, 35].

**Figure 2:**
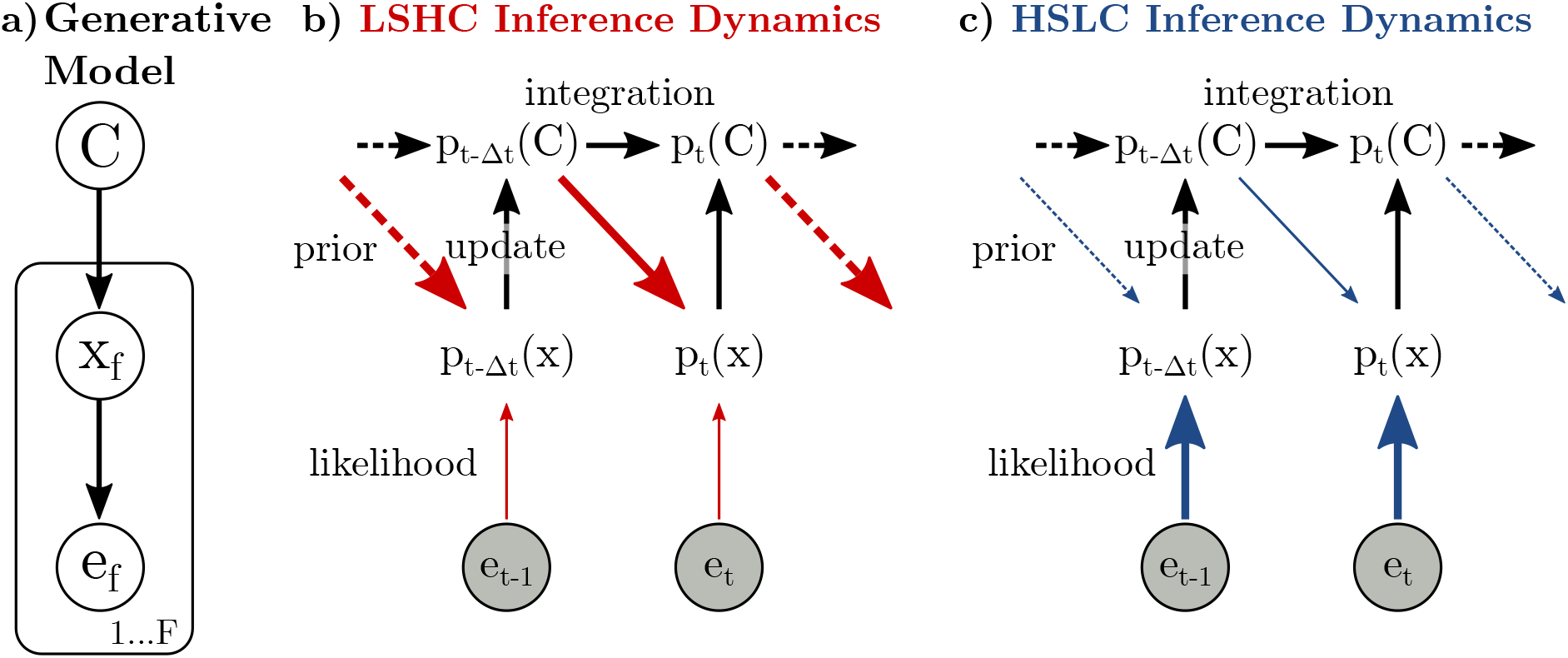
Information flow during hierarchical inference where categorical beliefs are fed back as a prior on sensory features. **a)** Generative model that we assume the brain has learned for a discrimination task, which specifies how sensory observations *e_f_* depend on the category for the trial, *C*, in two stages: each sensory observation *e_f_* is assumed to be a noisy realization of underlying sensory features, *x_f_*, and each frame of sensory features is itself assumed to be selected according to the trial’s category. **b-c)** Integrating evidence about *C* requires updating the current belief about *C* with new information derived from the sensory representation (left-right “integration” and bottom-up “update” arrows). The posterior distribution over *x* combines top-down expectations (diagonal “prior” arrows) with new evidence from the stimulus, *e_f_* (bottom-up “likelihood” arrows). Width of arrows indicates average amount of information communicated; red and blue arrows indicate changes in information flow between conditions. Note that when inference is exact, the prior is subtracted from the information in the update during the integration to prevent double-counting early evidence. While the generative model in (a) operates with discrete frames, *f*, inference in the brain happens in continuous time, *t*. **b)** LSHC: Low sensory information means little information in the likelihood about sensory features *x_f_*. High category information means that most of this information is also informative about *C*. It also means high information in the prior that is fed back to the sensory representation. **c)** HSLC: High sensory information means high information in the likelihood about sensory features *x_f_*. Low category information means that this information is only weakly predictive of *C*. It also means little information in the prior that is being fed back to the sensory representation.

We hypothesize that feedback of “decision-related” information to sensory areas [46, 16] implements a prior that reflects current beliefs about the stimulus category [27, 60, 35]. While such a prior is optimal from the perspective of estimating the sensory features, *x_f_*, this complicates evidence accumulation (Methods). When *x_f_* is influenced by prior beliefs about the stimulus category, the calculation of the “update” (log likelihood odds or LLO_*f*_) cannot simply replace p(*e_f_*|*C*) by p(*x_f_*|*C*); instead, the decision-making area now needs to account for or “divide out” the influence of the top-down prior on the sensory representation (Figure 2b-c). For an ideal observer performing exact inference, this process would not entail any suboptimalities or biases. However, inference in the brain is necessarily approximate, with the *potential* to induce a bias.

*Under*-correcting for this prior would lead to earlier frames entering into the update multiple times, giving them a larger weight in the final decision. Over multiple frames, the effect is a positive feedback loop between estimates of sensory features *x_f_* and the belief in *C*. This mechanism, which we call a “perceptual confirmation bias,” leads to primacy effects. *Over*-correcting for the prior, on the other hand, would lead to information from earlier frames decaying away, giving earlier frames less influence on the final decision and manifesting as recency effects. Importantly, in either case, the strength of any bias is directly related to the strength of the prior, which in turn is determined by the category information in the task (Figure 2).

We implemented two canonical models, corresponding to each of the two major classes of approximate inference schemes known from statistics and machine learning: sampling-based and variational inference [5, 40], and both of which have previously been proposed models for inference in the brain [20, 51, 55, 23]. In both cases, we assumed that sensory areas of the brain approximate the posterior, incorporating both the current sensory input *and* expectations based on past frames. Interestingly, both sampling-based and variational inference models behaved similarly in terms of performance and biases, and so here only show the results from the sampling-based model, and provide the corresponding variational results in the SI. The performance of our approximate models (Figure 3b) largely matched that of an exact inference model (Figure 3a), with accuracy somewhat reduced for high category information. We computed the temporal biases of the approximate inference models for each combination of sensory information and category information, and found that both models showed a primacy effect whose magnitude decreased with the amount of category information (Figure 3b-d). Both hierarchical inference models *under*-corrected for the influence of prior expectations on the sensory representation. Over the course of a trial, this lead to a positive feedback loop between the evidence-integration part of the model and the sensory representation, with the strength of this loop being strong in the LSHC and weak in the HSLC condition (Figure 2b-c). Importantly, this bias is a direct consequence of the *approximate* nature of the representation of the posterior distribution; for instance, in the sampling model, the bias disappears as the number of samples gets large (Methods).

**Figure 3:**
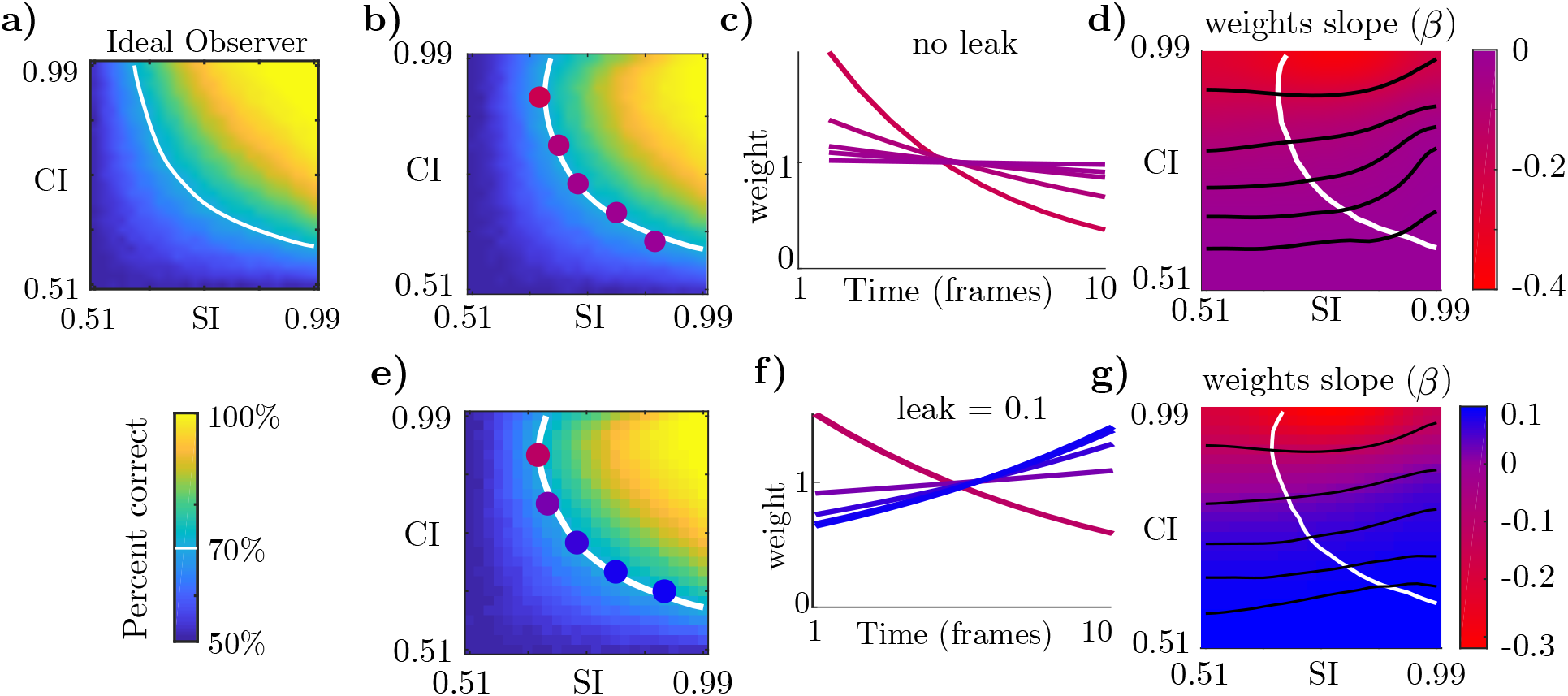
Changes in bias predicted by approximate hierarchical inference models. **a)** Performance of an ideal observer reporting *C* given ten frames of evidence. White line shows threshold performance, defined as 70% correct. The ideal observer’s temporal weights are always flat (not shown). **b)** Performance of our sampling-based approximate inference model with no leak (Methods). Colored dots correspond to lines in the next panel. **c)** Temporal weights in the model transition from flat to a strong primacy effect, all at threshold performance, as the stimulus transitions from the HSLC to the LSHC conditions. **d)** Visualization of how temporal biases change across the entire task space. Red corresponds to primacy, and blue to recency. White contour as in (b). Black lines are iso-contours for slopes corresponding to highlighted points in (b). **e-g)** Same as (b-d) but with leaky integration, which lessens primacy effects and produces recency effects when category information is low.

Results for the variational and sampling-based inference models are qualitatively the same (Supplemental Figure S4), as are results from simulating a larger neurally-inspired sampling model (Supplemental Figure S8) [27]. This indicates that the observed pattern of biases is not tied to a particular representation scheme – sampling or parametric – but to the *approximate* and *hierarchical* nature of inference.

Previous studies further suggest that evidence integration in the brain may be “leaky” or “forgetful,” which can be motivated either mechanistically [19, 13, 6, 64], or as an adaptation to non-stationary environments in normative models [25]. Including leaky integration, our final approximate inference models contain two competing mechanisms: first, they exhibit a confirmation bias as a consequence of approximate hierarchical inference, which is strongest when category information is high, leading to a primacy effect. Second, they contain leaky integration dynamics, which dampens the primacy effect and results in recency effects when category information is low and confirmation bias dynamics are weak (Figure 3e-g). While the exact magnitude of the leak is a free parameter in our model, to be constrained by data, the *change* in bias with changes in category information is a strong prediction, i.e. changing from strong primacy to no bias, or changing from weak primacy to recency, depending on the magnitude of the leak.

We performed additional simulations to explore the interaction between leaky integration and hierarchical inference. First, we observed that leaky integration can, surprisingly, *improve* performance, since it counteracts the confirmation bias when category information is high (Supplemental Figure S5). We further observed that optimizing the magnitude of the leak for maximum performance, separately for each combination of sensory information and category information, always resulted in flat temporal weights (Supplemental Figure S6).

Our models make a critical prediction that is not shared by any other model: that the temporal bias of the very same observer should change from primacy to flat or recency in a task in which nothing changes apart from the balance between category and sensory information.

### Changing category information in a visual discrimination task

To test this prediction, we designed an orientation discrimination task with two stimulus conditions that correspond to the two opposite sides of this task space (LSHC and HSLC), while keeping all other aspects of the design the same (Figure 4a-b). Importantly, in this experiment, within-observer comparisons between the two task conditions isolate relative changes in sensory information and category information. This overcomes the difficulties in directly quantifying sensory information and category information as “high” or “low” in an isolated task, which requires additional assumptions, as discussed above.

**Figure 4:**
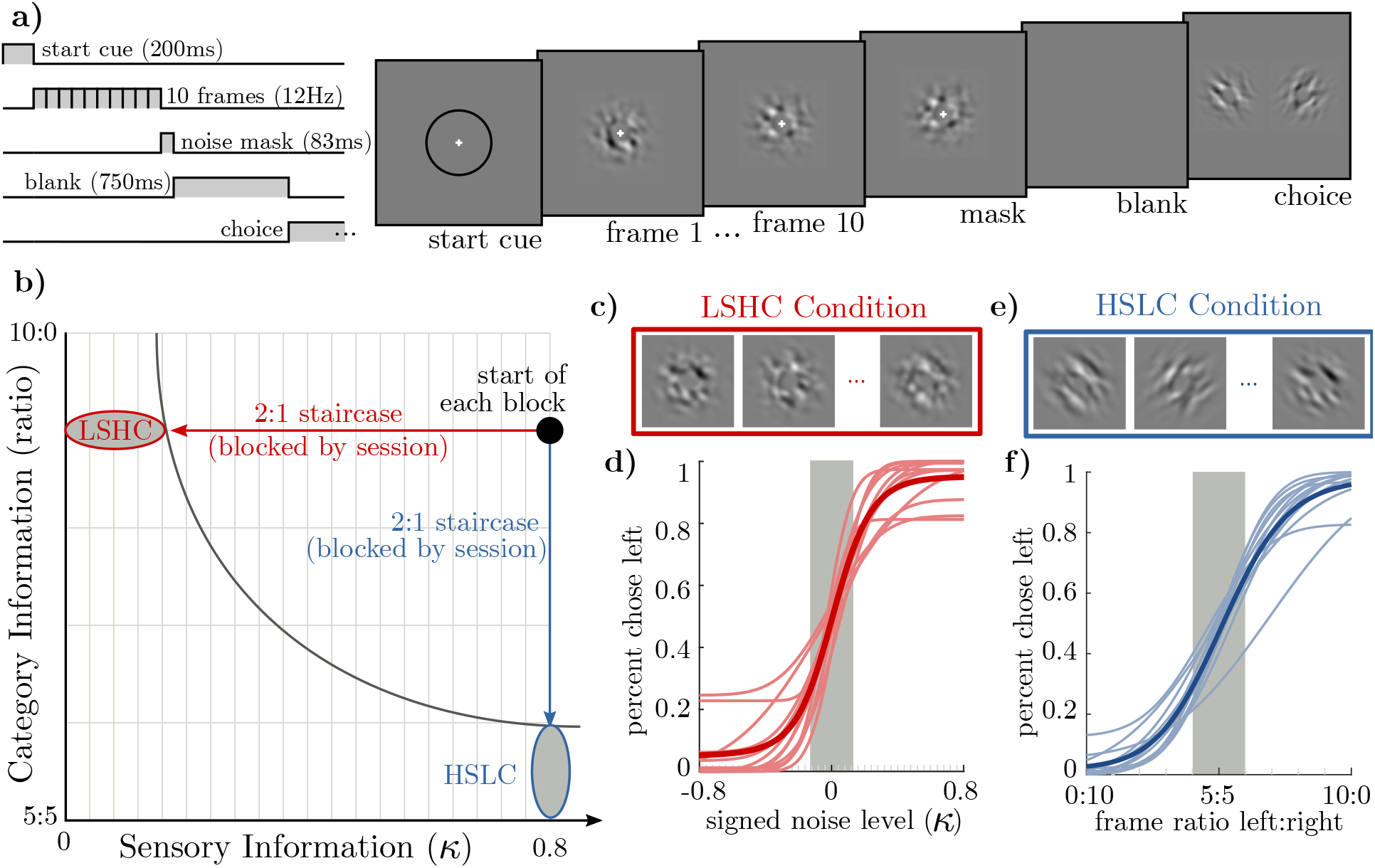
Two task conditions that reduce either sensory or category information to threshold level using a staircase. **a)** Each trial consisted of a 200ms start cue, followed by 10 stimulus frames presented for 83ms each, followed by a single mask frame of zero-coherence noise. After a 750ms delay, left or right targets appeared and participants pressed a button to categorize the stimulus as “left” or “right.” Stimulus contrast is amplified and spatial frequency reduced in this illustration. **b)** Category information is determined by the expected ratio of frames in which the orientation matches the correct category, and sensory information is determined by a parameter *κ* determining the degree of spatial orientation coherence (Methods). At the start of each block, we reset the staircase to the same point, with category information at 9 : 1 and *κ* at 0.8. We then ran a 2-to-1 staircase either on *κ* or on category information. The *Low-Sensory-High-Category (LSHC)* and *High-Category-Low-Sensory (HSLC)* ovals indicate sub-threshold trials; only these trials were used in the regression to infer observers’ temporal weights. **c)** Visualization of a noisy stimulus in the LSHC condition. All frames are oriented to the left. **d)** Psychometric curves for all observers (thin lines) and averaged (thick line) over the staircase. Shaded gray area indicates the median threshold level across all observers. **e)** Example frames in the HSLC condition. The orientation of each frame is clear, but orientations change from frame to frame. **f)** Psychometric curves over frame ratios, plotted as in (d).

The stimulus in our task consisted of a sequence of ten visual frames (Figure 4a). Each frame consisted of band-pass-filtered white noise with excess orientation power either in the 45° or the +45° orientation [1] (Figure 4b,d). On each trial, there was a single true orientation category, but individual frames might differ in their orientation. At the end of each trial, observers reported whether the stimulus was oriented predominantly in the −45° or the +45° orientation (Methods).

Sensory information in our task is determined by how well each image determines the orientation of that frame (i.e. the amount of “noise” in each frame), and category information is determined by the probability that any given frame’s orientation matches the trial’s category. We used signal detection theory to quantify both sensory information and category information as the area under the receiver-operating-characteristic curve for *e_f_* and *x_f_* (sensory information), and for *x_f_* and *C* (category information). Hence for a ratio of 5 : 5 frames of each orientation, a frame’s orientation does not predict the correct choice and category information is 0.5. For a ratio of 10 : 0, knowledge of the orientation of a single frame is sufficient to determine the correct choice and category information is 1. Quantifying sensory information depends on individual observer’s sensory noise, but likewise ranges from 0.5 to 1 (see Supplemental Text).

For each observer, we compared two conditions intended to probe the difference between the LSHC and HSLC regimes. Starting with a stimulus containing both high sensory and high category information, we either ran a 2:1 staircase lowering the sensory information while keeping category information high, or we ran a 2:1 staircase lowering category information while keeping sensory information high (Figure 4b). Sub-threshold trials in each condition define the LSHC and HSLC regimes, respectively (Figure 4c,e). For each condition and each observer, we used logistic regression to infer the influence of each frame onto their choice. observers’ overall performance was matched in the two conditions by setting a performance threshold below which trials were included in the analysis (Methods).

In agreement with our hypothesis, we find predominantly flat or decreasing temporal weights in the LSHC condition (Figure 5a), and when the information is partitioned differently – in the HSLC condition – we find flat or increasing weights (Figure 5b). To quantify this change, we first used cross-validation to select a method for quantifying temporal slopes, and found that constraining weights to be either a linear or exponential function of time worked equally well, and both outperformed logistic regression with a smoothness prior (Supplemental Figure S2d; Methods). A within-observer comparison revealed that the change in slope between the two conditions was as predicted for all observers (Figure 5h) (*p* < 0.05 for 9 of 12 observers, bootstrap). The effect was also highly significant on a population level (*p* < 0.01, sign test on median slope parameters for each observer). This demonstrates that the trade-off between sensory and category information in a task robustly changes observers’ temporal weighting strategy as we predicted.

**Figure 5:**
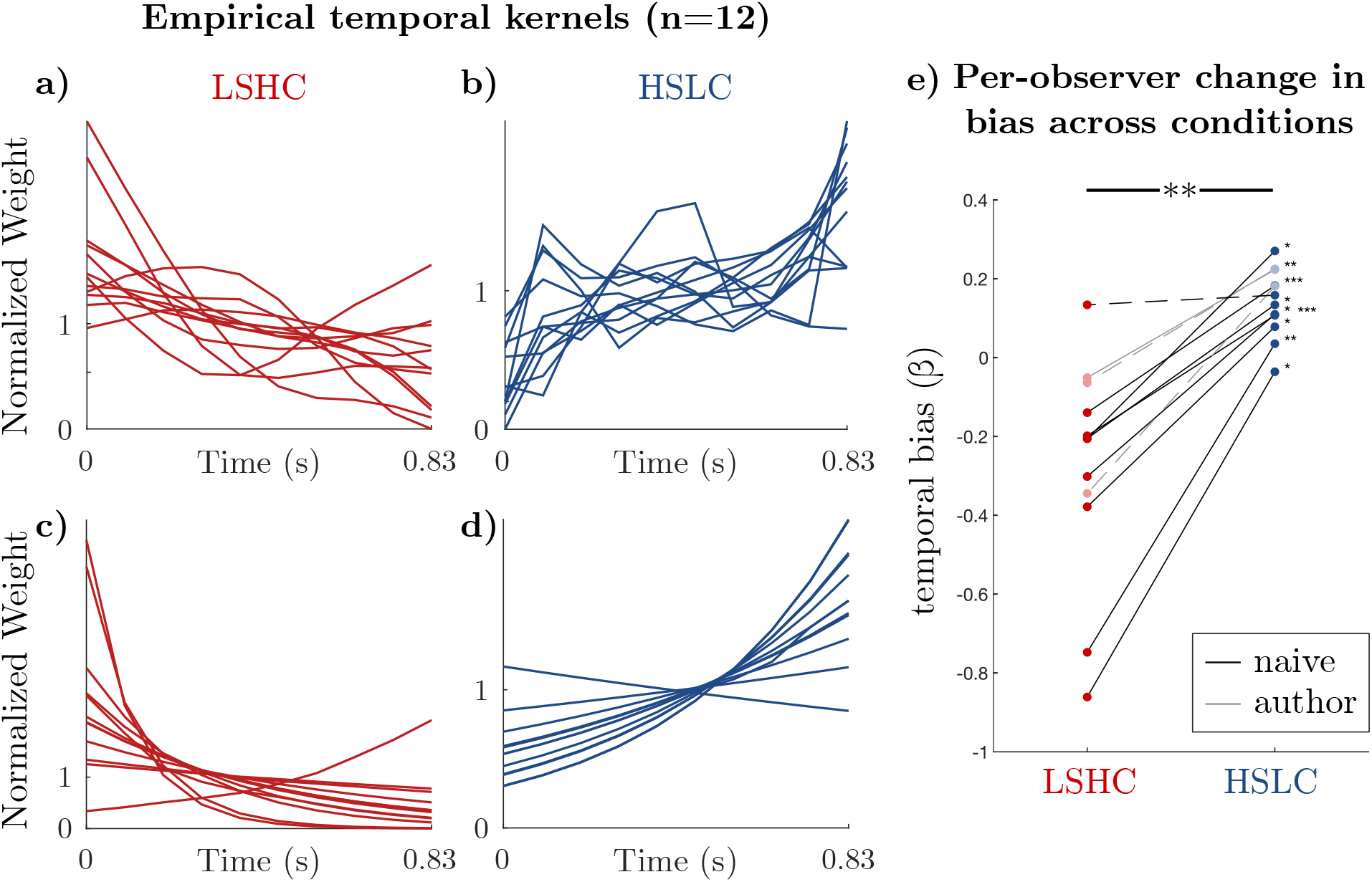
Every observer’s temporal bias consistently changes from primacy to unbiased/recency between conditions as predicted. **a-b)** Temporal weights from logistic regression of choices from sub-threshold frames for individual observers. Weights are regularized by a cross-validated smoothness term, and are normalized to have a mean of 1. **c-d)** To summarize temporal biases, we constrained weights to be an exponential function of time and re-fit them to observers’ choices. Exponential weights had higher cross-validated performance than regularized logistic regression, supporting their use to summarize temporal biases (Supplemental Figure S2d; Methods). **e)** The *change* in the temporal bias, quantified as the exponential slope parameter (*β*), between the two task contexts for each observer is consistently positive (combined, *p* < 0.01, sign test on median slope from bootstrapping). This result is individually significant in 9 of 12 observers by bootstrapping (*p* < 0.05, *p* < 0.01, and *p* < 0.001 indicated by *, **, and ** * respectively; non-significant observers plotted with dashed lines). Points are median slope values after bootstrap-resampling each observer’s sub-threshold trials. A slope parameter *β* > 0 corresponds to a recency bias and *β* < 0 to a primacy bias. We found similar results using linear rather than exponential weight functions (Supplemental Figure S3).

### Confirmation bias, not bounded integration, explains primacy effects

The primary alternative explanation for primacy effects in fixed-duration integration tasks proposes that observers integrate evidence to an internal *bound*, at which point they cease paying attention to the stimulus [33]. In this scenario, early evidence always enters the decision-making process while evidence late in trial is often ignored. Averaged over many trials, this results in early evidence having a larger effect on the final decision than late evidence, and hence decreasing regression weights (and psychophysical kernels) just as we found in the LSHC condition. Both models reflect very different underlying mechanism: in our approximate hierarchical inference models, a confirmation bias ensures that early evidence has a larger effect on the final decision than late evidence for every single trial. In the integration-to-bound (ITB) model, in a single trial, all evidence is weighed exactly the same before the bound is reached, and not at all afterwards.

In order to test whether the ITB mechanism or confirmation bias dynamics better explain our data, we used an Extended ITB model that can account for both biased integration dynamics (as during a confirmation bias), and for incomplete evidence accumulation due to a finite bound [31]. This model is a simple extension to classic drift diffusion models [26]. Until it hits a bound or the trial ends, the model integrates signals as follows:

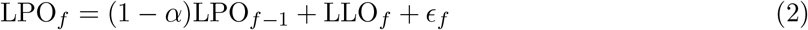

where *∈_f_* represents noise in the accumulation process (Figure 6a, Methods). For *α* = 0, this model weighs evidence equally (optimally) over time. *α* > 0 has previously been proposed to model “forgetful integration” in mechanistic models [13, 6, 26, 11], or as reflecting an assumption of non-stationarity in the environment in normative models [25], and leads to a recency effect. Importantly, a negative value for *α* induces “accelerating” integration dynamics, in which already-accumulated evidence is amplified, leading to primacy effects [13, 6, 31].

**Figure 6:**
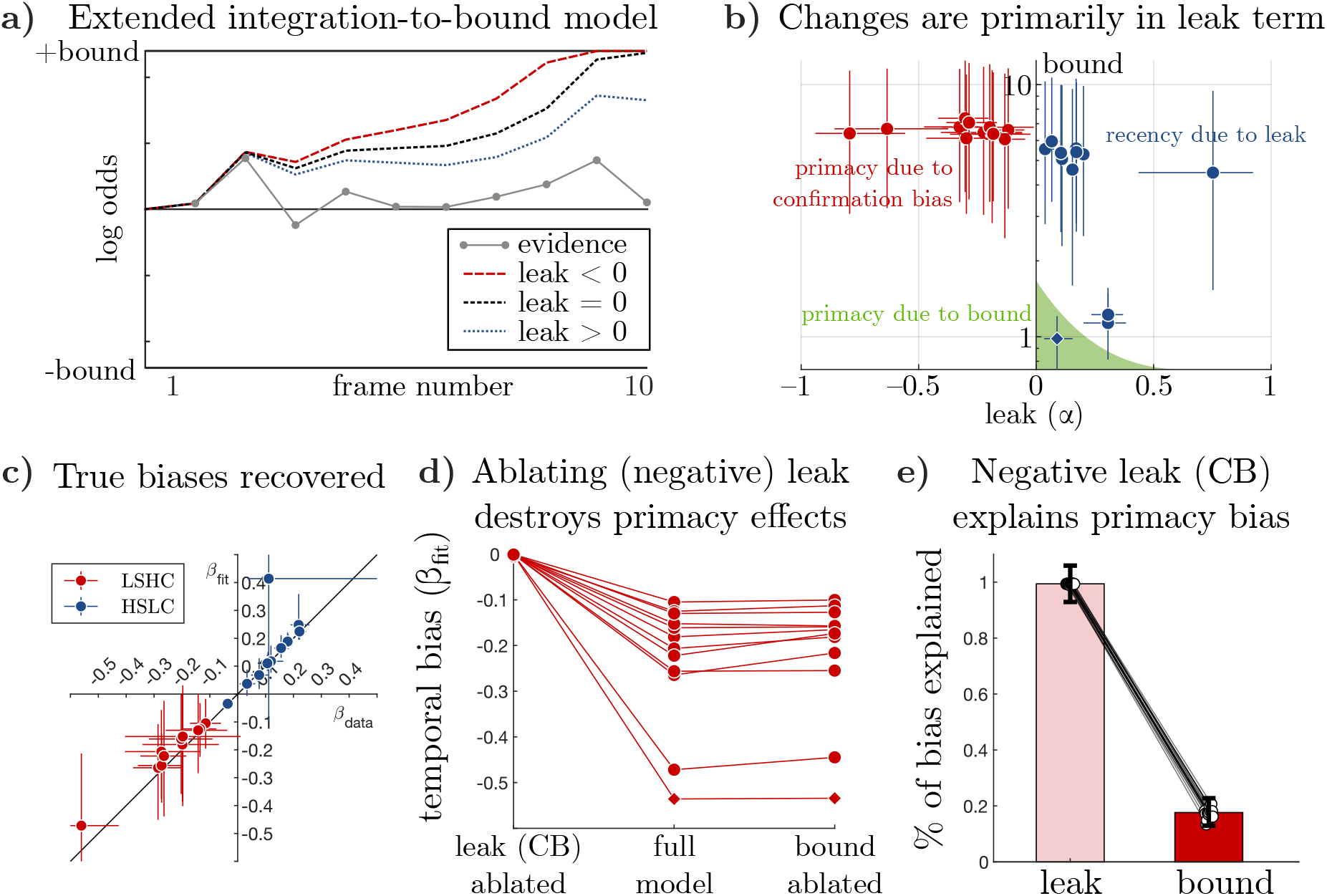
Fitting an Extended Integration-to-Bound (“Extended ITB”) model to data demonstrates that integration dynamics (negative ff for confirmation bias, positive ff for forgetting), rather than a bound, best accounts for data. **a)** Illustration of Extended ITB model. As in classic drift-diffusion models with an absorbing bound, evidence is integrated to an internal bound, after which new evidence is ignored. Compared to perfect integration (*α* = 0), a *positive* leak (*α* > 0) decays information away and results in a recency bias, and a negative leak (*α* < 0) amplifies already integrated information, resulting in a primacy bias. Since *α* < 0 may also result in more bound crossings, both leak and bound together determine the shape of the temporal weights. **b)** Inferred values of the bound and leak parameters in each condition, shown as median±68% credible intervals. The classic ITB explanation of primacy effects corresponds to a non-negative leak and a small bound – illustrated here as a shaded green area. Note that the three observers near the ITB regime are points from the HSLC task – two still exhibit mild recency effects and one exhibits a mild primacy effect as predicted by ITB. **c)** Across both conditions, the temporal slopes (*β*) implied by the full model fits closely match the slopes in the data. *β* < 0 corresponds to primacy, and *β* > 0 to recency. Error bars indicate 68% confidence intervals from bootstrapping trials on *β*_data_ and from posterior samples on *β*_fit_. **d)** Median temporal biases implied by the full model (middle) and by the model with either zero leak (left) or infinite bound (right). Each line corresponds to a single observer. (LSHC condition only – HSLC condition in Supplemental Figure S12). d) Across the population, the negative leak (confirmation bias) accounted for 99% (68%CI=[93%, 106%]), and bounded integration accounted for 18% (68%CI=[13%, 23%]) of the primacy bias captured by the model. (Additional analyses in Supplemental Figure S12).

The Extended ITB model produces three distinct patterns in the data (colored text in Figure 6c). First, when *α* is positive and the bound is large, it produces recency biases. Second, when the bound is small, it produces primacy biases [33], as long as *α* is also small so that it does not prevent the bound from being crossed. Third, when the bound is large and *α* is *negative*, it also produces primacy biases but now due to confirmation-bias-like dynamics rather than due to bounded integration. Crucially, this single model can account for both primacy due to a bound and primacy due to a confirmation bias by different parameter values (recovery of ground-truth mechanisms shown in Supplemental Figures S10, S11). Examining the parameters of this model fit to data therefore allows us to determine the relative contributions of bounded integration and confirmation bias dynamics in cases where observers show primacy effects.

We fit the Extended ITB model to individual choices on sub-threshold trials, separately for the LSHC and the HSLC conditions. Figure 6c shows the posterior mean and 68% credible interval for the dynamics parameter, ff, and bound parameter inferred for each observer. The model consistently inferred a negative *α* in the LSHC condition and a positive *α* in the HSLC condition for all observers, suggesting that confirmation-bias dynamics are crucial to explain observer’s primacy biases in the LSHC condition, as well as the change in bias from LSHC to HSLC conditions. However, while the inferred bound for every single observer is so high as not to contribute at all *if the leak was zero*, it is possible that bounded integration still contributes to primacy effects, given that a stronger confirmation bias (*α* < 0) will hit a bound more often.

We therefore performed an ablation analysis to quantify the relative contribution of the leak and bound parameters to the primacy effect in the LSHC condition (Methods). We first asked whether the inferred model parameters reproduced the observed biases. Indeed, Figure 6b shows near-perfect agreement between the temporal biases implied by simulating choices from the fitted models and the biases inferred directly from observers’ choices. Given this, if setting the bound to 1 leaves temporal biases unchanged, then we can conclude that biases were driven by the leak, and conversely, a temporal bias that remains after setting *α* to zero must be due to the bound. Figure 6d shows that primacy effects largely disappear when *α* is ablated, but not when the bound is ablated. To summarize ablation effects across the population, we used a hierarchical regression model to compute a population-level “ablation index” for each parameter, which is 0 if removing the parameter has no effect on temporal slopes, *β*, and is 1 if removing it destroys all temporal biases (Methods). The ablation index can therefore be interpreted as the fraction of the observers’ primacy or recency biases that are attributable to each parameter (but they do not necessarily sum to 1 because the slope is a nonlinear combination of both parameters). In the LSHC condition, the ablation index for the leak term was 0.99 (68% CI=[0.93, 1.06]), and for the bound term it was 0.18 (68% CI=[0.13, 0.23]) (Figure 6e). This indicates that although both mechanisms are present, primacy effects in our data are dominated by the self-reinforcing dynamics of a negative leak. Results for the HSLC condition are shown in Supplemental Figure S12.

Interestingly, one observer exhibited a slight primacy effect in the HSLC condition, and our analyses suggest this was primarily due to bounded integration dynamics as proposed by Kiani et al (2008). This outlier observer is marked with a diamond symbol throughout Figure 6, and is further highlighted in Supplemental Figure S12. However, even this observer’s primacy effect in the LSHC condition was driven by a confirmation bias (negative ff), and their change in slope between LSHC and HSLC conditions was in the same direction as the other observers. Importantly, finding a primacy effect due to an internal bound confirms that our model fitting procedure is able to detect such effects when they are, in fact, present. Two additional observers appear to have low bounds in the HSLC condition (Figure 6c), but are dominated by leaky integration (*α* > 0), resulting in an overall recency bias.

## Discussion

Our work makes three main contributions. First, we extended ideal observer models of evidence integration tasks by explicitly accounting for the intermediate sensory representation. We showed that this partitions the information in the stimulus about the category into two parts – “sensory information” and “category information” – defining a novel two-dimensional space of possible tasks. Second, we found that two classes of biologically-plausible approximate inference algorithms entailed a confirmation bias whose strength strongly varied across this task space. Interestingly, the location of tasks in existing studies qualitatively predicted the bias they found across species, sensory modalities and task designs. Third, we collected new data and confirmed a critical prediction of our theory, namely that individual observers’ temporal biases should change depending on the balance of sensory information and category information in the stimulus. Finally, by fitting an extended integration to bound (Extended ITB) model to individual observer choices, we confirmed that these changes in biases are due to a change in integration dynamics rather than bounded integration.

The “confirmation bias” emerges in our hierarchical inference models as the result of three key assumptions. Our first assumption is that inference in evidence integration tasks is in fact hierarchical, and that the brain approximates the posterior distribution over the intermediate sensory variables, *x*. This is in line with converging evidence that populations of sensory neurons encode posterior distributions of corresponding sensory variables [36, 78, 4, 2] incorporating dynamic prior beliefs via feedback connections [36, 78, 2, 45, 60, 47, 27, 35]. This is in contrast to other probabilistic theories in which only the likelihood is represented in sensory areas [38, 3, 48, 69], which would not predict primacy effects due to confirmation bias dynamics.

Our second key assumption is that evidence is accumulated online. In our models, the belief over *C* is updated based only on the posterior from the previous step and the current posterior over *x*. This can be thought of as an assumption that the brain does not have a mechanism to store and retrieve earlier frames of evidence directly, and is consistent with drift-diffusion models of decision-making [26]. As mentioned in the main text, the assumptions until now – hierarchical inference with online updates – do not entail any temporal biases for an ideal observer. Further, the use of discrete time in our experiment and models is only for mathematical convenience – analogous dynamics emerge in continuous-time, and in fact we implemented our models at a finer time scale than at which evidence frames are presented.

Third, we assumed that inference in the brain is approximate – a safe assumption due to the intractable nature of exact inference in large models. In the sampling model, we assumed that the brain can draw a limited number of independent samples of *x* per update to *C*, and found that for a finite number of samples the model is inherently unable to account for all of the prior bias of *C* on *x* in its updates to *C*. Existing neural models of sampling typically assume that samples are distributed temporally [29, 20, 12, 47, 27], but it has also been proposed that the brain could process multiple sampling “chains” distributed spatially [56]. The relevant quantity for our model is the number of independent samples that can be tallied per update: the more samples, the smaller the bias. The variational model’s representational capacity was limited by enforcing that the posterior over *x* is unimodal, and that there is no explicit representation of dependencies between *x* and *C*. Importantly, this does not imply that *x* and *C* do not influence each other. Rather, the Variational Bayes algorithm expresses these dependencies in the *dynamics* between the two areas: each update that makes *C* = +1 more likely pushes the distribution over *x* further towards +1, and vice versa. Because the number of dependencies between variables grows exponentially, such approximates are necessary in variational inference with many variables [20]. The Mean Field Variational Bayes algorithm that we use here has been previously proposed as a candidate algorithm for neural inference [53].

The assumptions up to now predict a primacy effect but cannot account for the observed recency effects. When we incorporate a forgetting term in our models, they reproduce the observed range of biases from primacy to recency. The existence of such a forgetting term is supported by previous literature [67, 6]. Further, it is normative in our framework in the sense that reducing the bias in the above models improves performance (Supplemental Figures S4-S6). The optimal amount of bias correction depends on the task statistics: for high category information where the confirmation bias is strongest, a stronger forgetting term is needed to correct for it. While it is conceivable that the brain would optimize this term to the task [11, 50], our data suggest it is stable across our LSHC and HSLC conditions, or only adapts slowly.

It has been proposed that post-decision feedback biases subsequent perceptual estimations [59, 62]. While in spirit similar to our confirmation bias model, there are two conceptual differences between these models and our own: First, the feedback from decision area to sensory area in our model is both continuous and online, rather than conditioned on a single choice after a decision is made. Second, our models are derived from an ideal observer and only incur bias due to algorithmic approximations, while previously proposed “self-consistency” biases are not normative and require separate justification.

Our analysis decisively shows that accelerating dynamics, rather than reaching a bound before the end of the trial, explains the primacy effect in our data. Prior work has suggested that such accelerating dynamics may arise from two attractor states corresponding to the two choices *within* a decision-area [70, 74, 75, 71, 73], and that the nature of the temporal bias depends on the volatility of the integrated signal [52]. Importantly, decision-dynamics alone *cannot* explain our results, since the input to the attractors in such models usually reflects the total information in each frame about the choice, i.e. combining both sensory and category information. In other words, attractor models usually integrate *log odds*, which we kept approximately constant between LSHC and HSLC conditions. The same argument applies to other models that do no distinguish between sensory and category information, whether based on mixing trials of different difficulty [17] or differential accumulation of consistent and inconsistent evidence [62, 66, 34, 63].

In contrast, in our explanation based on approximate hierarchical inference, attractor dynamics arise *across* sensory and decision areas, as the result of cortical inter-area feedback whose strength is monotonically related to category information. Holding the task difficulty (and hence the magnitude of the log odds of each frame) constant, our model nonetheless predicts stronger inter-area attractor dynamics when category information is high. Given recent evidence that noise correlations contain a task-dependent feedback component [7], we therefore predict a reduction of task-dependent noise correlations in comparable tasks with lower category information. The confirmation bias mechanism may also account for the recent finding that stronger attractor dynamics are seen in a categorization task than in a comparable estimation task [61].

It has also been proposed that primacy effects could be the result of near-perfect integration of an adapting sensory population [73, 77]. For this mechanism to explain our full results, however, the sensory population would need to become less adapted over the course of a trial in our HSLC condition, while at the same time *more* adapted in the LSHC condition. We are unaware of such an adaptation mechanism in the literature. Further, such stimulus-dependent circuit dynamics would not predict top-down neural effects such as the task-dependence of the dynamics of sensory populations [61] nor the origin and prevalence of differential correlations [7], both of which are consistent with hierarchical inference [27, 35].

While our focus is on the perceptual domain in which observers integrate evidence over a timescale on the order of tens or hundreds of milliseconds, analogous computational principles hold in the cognitive domain over longer timescales. The crucial computational motif underlying our model of the confirmation bias is approximate hierarchical inference over multiple timescales. An agent in such a setting must simultaneously make accurate judgments of current data (based on the current posterior) and track long-term trends (based on all likelihoods). For instance, Zylberberg et al. (2018) identified an analogous challenge when observers must simultaneously make categorical decisions each trial (their “fast” timescale) while tracking the stationary statistics of a block of trials (their “slow” timescale), with trial-by-trial statistics analogous to the frame-by-frame statistics in our LSHC condition. As the authors describe, if observers base model updates on posteriors rather than likelihoods, they will further entrench existing beliefs [79]. However, the authors did not investigate order effects; our proposed confirmation bias models would predict that observers’ estimates of block statistics is biased towards earlier trials in the block (primacy). Schustek et al. (2018) likewise asked observers to track information across trials in a cognitive task more analogous to our HSLC condition, and report close to flat weighting of evidence across trials [57] in agreement with our model.

The strength of the perceptual confirmation bias is directly related to the integration of internal “top-down” beliefs and external “bottom-up” evidence previously implicated in clinical dysfunctions of perception [30]. Therefore, the differential effect of sensory and category information may be useful in diagnosing clinical conditions that have been hypothesized to be related to abnormal integration of sensory information with internal expectations [21].

Hierarchical (approximate) inference on multiple timescales is a common motif across perception, cognition, and machine learning. We suspect that all of these areas will benefit from the insights on the causes of the confirmation bias mechanism that we have described here and how they depend on the statistics of the inputs in a task.

## Methods

### Visual Discrimination Task

We recruited students at the University of Rochester as observers in our study. All were compensated for their time, and methods were approved by the Research observers Review Board. We found no difference between naive observers and authors, so all main-text analyses are combined, with data points belonging to authors and naive observers indicated in Figure 5d.

Our stimulus consisted of ten frames of band-pass filtered noise [1, 44] masked by a soft-edged annulus, leaving a “hole” in the center for a small cross on which observers fixated. The stimulus subtended 2.6 degrees of visual angle around fixation. Stimuli were presented using Matlab and Psychtoolbox on a 1920×1080px 120 Hz monitor with gamma-corrected luminance [8]. Observers kept a constant viewing distance of 36 inches using a chin-rest. Each trial began with a 200ms “start” cue consisting of a black ring around the location of the upcoming stimulus. Each frame lasted 83.3ms (12 frames per second). The last frame was followed by a single double-contrast noise mask with no orientation energy. Observers then had a maximum of 1s to respond, or the trial was discarded (Supplemental Figure 4)a. The stimulus was designed to minimize the effects of small fixational eye movements: (i) small eye movements do not provide more information about either orientation, and (ii) each 83ms frame was too fast for observers to make multiple fixations on a single frame.

The stimulus was constructed from white noise that was then masked by a kernel in the Fourier domain to include energy at a range of orientations and spatial frequencies but random phases [1, 44, 7] (a complete description and parameters can be found in the Supplemental Text). We manipulated sensory information by broadening or narrowing the distribution of orientations present in each frame, centered on either +45° or −45° depending on the chosen orientation of each frame. We manipulated category information by changing the proportion of frames that matched the orientation chosen for that trial. The range of spatial frequencies was kept constant for all observers and in all conditions.

Trials were presented in blocks of 100, with typically 8 blocks per session (about 1 hour). Each session consisted of blocks of only HSLC or only LSHC trials (Figure 4). Observers completed between 1500 and 4400 trials in the LSHC condition, and between 1500 and 3200 trials in the HSLC condition. After each block, observers were given an optional break and the staircase was reset to *κ* = 0.8 and *p*_match_ = 0.9. *p*_match_ is defined as the probability that a single frame matched the category for a given trial. In each condition, psychometric curves were fit to the concatenation of all trials from all sessions using the Psignifit Matlab package [58], and temporal weights were fit to all trials below each observer’s threshold.

### Low Sensory-, High Category-Information (LSHC) Condition

In the LSHC condition, a continuous 2-to-1 staircase on *κ* was used to keep observers near threshold (*κ* was incremented after each incorrect response, and decremented after two correct responses in a row). *p*_match_ was fixed to 0.9. On average, observers had a threshold (defined as 70% correct) of *κ* = 0.17±0.07 (1 standard deviation). Regression of temporal weights was done on all sub-threshold trials, defined per-observer.

### High Sensory-, Low Category-Information (HSLC) Condition

In the HSLC condition, the staircase acted on *p*_match_ while keeping fixed at 0.8. Although *p*_match_ is a continuous parameter, observers always saw 10 discrete frames, hence the true ratio of frames ranged from 5:5 to 10:0 on any given trial. Observers were on average 69.5% ± 4.7% (1 standard deviation) correct when the ratio of frame types was 6:4, after adjusting for individual biases in the 5:5 case. Regression of temporal weights was done on all 6:4 and 5:5 ratio trials for all observers, regardless of the underlying *p*_match_ parameter.

### Logistic Regression of Temporal Weights

We constructed a matrix of per-frame signal strengths S on sub-threshold trials by measuring the empirical signal level in each frame. This was done by taking the dot product of the Fourier-domain energy of each frame as it was displayed on the screen (that is, including the annulus mask applied in pixel space) with a difference of Fourier-domain kernels at +45° and 45° with = 0.16. This gives a scalar value per frame that is positive when the stimulus contained more +45° energy and negative when it contained more −45° energy. Signals were z-scored before performing logistic regression, and weights were normalized to have a mean of 1 after fitting.

Temporal weights were first fit using (regularized) logistic regression with different types of regularization. The first regularization method consisted of an AR0 (ridge) prior, and an AR2 (curvature penalty) prior. We did not use an AR1 prior to avoid any bias in the slopes, which is central to our analysis.

To visualize regularized weights in Figure 5, the ridge and AR2 hyperparameters were chosen using 10-fold cross-validation for each observer, then averaging the optimal hyperparameters across observers for each task condition. This cross validation procedure was used only for display purposes for individual observers in Figure 5a-b of the main text, while the linear and exponential fits (described below) were used for Figure 5c-d and statistical comparisons. Supplemental Figure S1 shows individual observers’ weights for all regression models.

We used two methods to quantify the shape (or slope) of **w**: by constraining **w** to be either an exponential or linear function of time, but otherwise optimizing the same maximum-likelihood objective as logistic regression. Cross-validation suggests that both of these methods perform similarly to either unregularized or the regularized logistic regression defined above, with insignificant differences (Supplemental Figure S2). The exponential is defined as

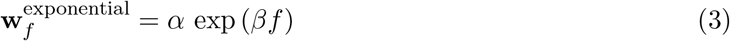

where *f* refers to the frame number. *β* gives an estimate of the shape of the weights w over time, while *α* controls the overall magnitude. *β* > 0 corresponds to recency and *β* < 0 to primacy. The *β* parameter is reported for human observers in Figure 5d, and for the models in Figure 3e,h.

The second method to quantify slope was to constrain the weights to be a linear function in time:

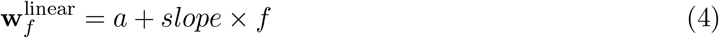

where *slope* > 0 corresponds to recency and *slope* < 0 to primacy.

Figure 5d shows the median exponential shape parameter (*β*) after bootstrapped resampling of trials 500 times for each observer. Both the exponential and linear weights give comparable results (Supplemental Figure S3).

Because we are not explicitly interested in the magnitude of **w** but rather its *shape* over stimulus frames, we always plot a “normalized” weight, **w**/mean(**w**), both for our experimental results (Figure 5a-d) and for the model (Figure 3c, f).

### Approximate inference models

We model evidence integration as Bayesian inference in a three-variable generative model (Figure 2a) that distills the key features of online evidence integration in a hierarchical model [27]. The variables in the model are mapped onto the sensory periphery (*e*), sensory cortex (*x*), and a decision-making area (*C*) in the brain. For simulations, the same model was used both to generate data (*C* → *x_f_* → *e_f_*), and, in the reverse direction, as a model of inference dynamics (*e_f_* → p(*x_f_* | …) ↔ p(*C*| …)).

In the generative direction, on each trial, the binary value of the correct choice *C* ∈ {−1, +1} is drawn from a 50/50 prior. *x_f_* is then drawn from a mixture of two Gaussians:

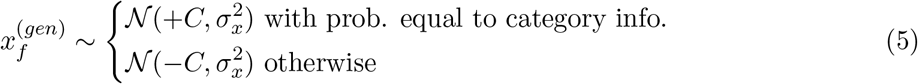

Finally, each *e_f_* is drawn from a Gaussian around *x_f_*:

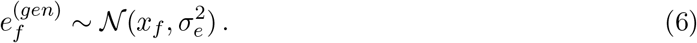

In the inference direction, we assume that the observer has learned the correct model parameters (namely the category information, and sensory information or 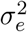), even as parameters change between the two different conditions. This is why we ran our observers in blocks of only LSHC or HSLC trials on a given day.

Category information in this model can be quantified by the probability that 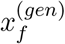 is drawn from the mode that matches *C*, as in equation (5). We quantify sensory information as the probability with which an ideal observer can recover the sign of *x_f_* from a single *e_f_*. That is, in our model sensory information is equivalent to the area under the ROC curve for two univariate Gaussian distributions separated by a distance of 2, which is given by

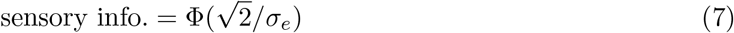

where ϕ is the inverse cumulative normal distribution.

Optimal inference of *C* requires conditioning on all frames of evidence *e*_1_,…, *e_f_*, which can be expressed as the Log Posterior Odds (LPO),

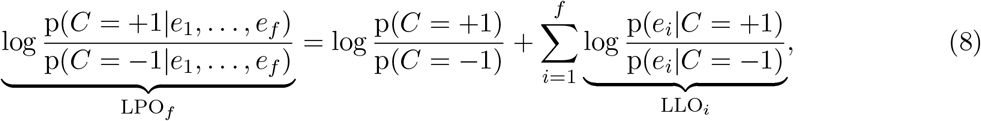

where LLO_*f*_ is the log likelihood odds for frame *f* [26, 6]. To reflect the fact that the brain has access to only one frame of evidence at a time, this can be rewritten this as an *online* update rule, summing the previous frame’s log posterior with new evidence gleaned on the current frame:

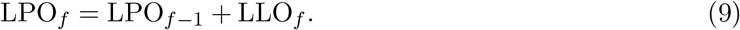

Optimal inference of *x_f_* similarly requires accounting for all possible sources of information. Ideally, sensory areas should incorporate prior information based on previous frames to compute p(*x_f_* |*e*_1_,…, *e_f_*). Using p_*f*-1_(*C* = *C*) = p(*C* = *c*|*e*_1_,…, *e*_*f*-1_) to denote the brain’s belief that the category is *C* = *c* after the first *f* − 1 frames, the posterior over *x_f_* given all frames, p(*x_f_*|*e*_1_,…, *e_f_*), can be written as

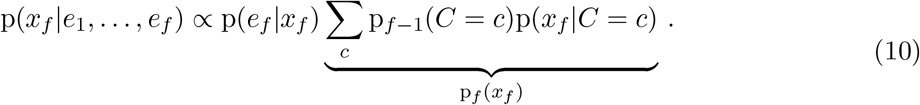

The term p_*f*_ (*x_f_*) is a prior on sensory features *x_f_* that changes over time depending on the current belief in the category, p_*f*-1_(*C*). In other words, sensory areas could dynamically combine instantaneous evidence (p(*e_f_*|*x_f_*)) with accumulated categorical beliefs (p_*f*-1_(*C*)) to arrive at a more precise estimate of present sensory features *x_f_*. This is what we mean by feedback of “decision-related information” or of “categorical beliefs.”

Our approximate inference models, described in detail below, compute a biased estimate of LLO_*f*_, which we call 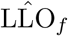. The bias is due to the interaction of approximations with feedback of prior beliefs, such that 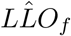 is biased towards LPO_*f*-1_, resulting in a confirmation bias. Importantly, this bias arises naturally in both the sampling-based and variational approximate inference algorithms that we study here, as a direct consequence the *approximate* nature of the posterior. Given the approximate representation of posteriors over *x_f_*, there is no way to exactly divide out the influence of the prior and recover the exact log likelihood odds on a frame-by-frame basis. However, the confirmation bias can be mitigated on average simply by incorporating a leak term, *γ*, in the integration process [13, 6]:

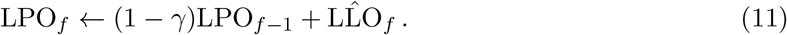

Due to the bias in 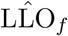, *γ* can be seen as a kind of approximate *bias correction*, with positive values for *γ* often improving performance (Supplemental Figure S5–S6). Equivalently, one can view the quantity 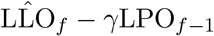 as a less biased estimate of the true log likelihood odds.

Because the effective time per update in the brain is likely faster than our 83ms stimulus frames, we included an additional parameter *n*_U_ for the number of online belief updates per stimulus frame. In the sampling model described below, we amortize the per-frame updates over *n*_U_ steps, updating *n*_U_ times per frame using 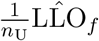. In the variational model, we interpret *n*_U_ as the number of coordinate ascent steps per stimulus frame.

Simulations of both models were done with 10000 trials per task type and 10 frames per trial. To quantify the evidence-weighting of each model, we used the same logistic regression procedure that was used to analyze human observers’ behavior. In particular, temporal weights in the model are best described by the exponential weights (equation (3)), so we use *β* to characterize the model’s biases.

### Sampling model

The sampling model estimates p(*e_f_*|*C*) using importance sampling of *x*, where each sample is drawn from a pseudo-posterior using the current running estimate of *p*_*f*-1_(*C*) ≡ *p*(*C*|*e*_1_, .., *e*_*f*-1_) as a marginal prior:

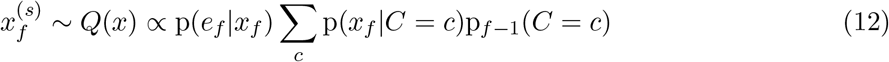

Using this distribution, we obtain the following unnormalized importance weights.

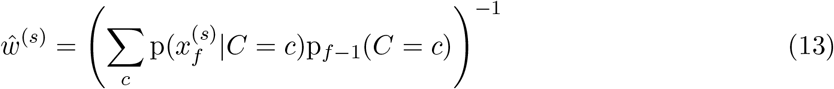

In the self-normalized importance sampling algorithm these weights are then normalized as follows,

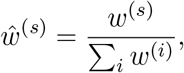

though we found that this had no qualitative effect on the model’s ability to reproduce the trends in the data. The above equations yield the following estimate for the log-likelihood ratio needed for the belief update rule in equation (11):

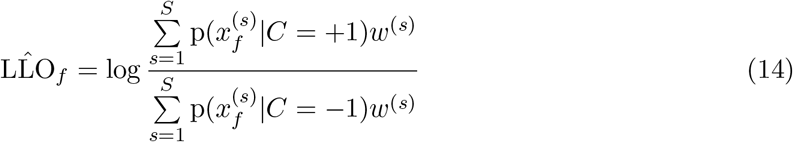

In the case of infinitely many samples, these importance weights exactly counteract the bias introduced by sampling from the posterior rather than likelihood, thereby avoiding any double-counting of the prior, and hence, any confirmation bias [49]. However, in the case of finite samples, *S*, biased estimates of LLO_f_ are unavoidable [15].

The full sampling model is given in Supplemental Algorithm S1. Simulations in the main text were done with *S* = 5, *n*_U_ = 5, normalized importance weights, and *γ* = 0 or *γ* = 0.1.

### Variational model

The following variational model produces qualitatively similar patterns of temporal biases to the IS model (Supplemental Figure S4).

The core assumption of the variational model is that while a decision area approximates the posterior over *C* and a sensory area approximates the posterior over *x*, no brain area explicitly represents posterior dependencies between them. That is, we assume the brain employs a *mean field approximation* to the joint posterior by factorizing p(*C*, *x*_1_,…, *x_F_*|*e*_1_,…, *e_F_*) into a product of approximate marginal distributions 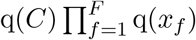 and minimizes the Kullback-Leibler divergence between q and p using a process that can be modeled by the Mean-Field Variational Bayes algorithm [40].

By restricting the updates to be online (one frame at a time, in order), this model can be seen as an instance of “Streaming Variational Bayes” [9]. That is, the model computes a sequence of approximate posteriors over *C* using the same update rule for each frame. We thus only need to derive the update rules for a single frame and a given prior over *C*; this is extended to multiple frames by re-using the posterior from frame *f* - 1 as the prior on frame *f*.

As in the sampling model, this model is unable to completely discount the added prior over *x*. Intuitively, since the mean-field assumption removes explicit correlations between *x* and *C*, the model is forced to commit to a marginal posterior in favor of *C* = +1 or *C* = −1 and *x* > 0 or *x* < 0 after each update, which then biases subsequent judgments of each.

To keep conditional distributions in the exponential family (which is only a matter of mathematical convenience and has no effect on the ideal observer), we introduce an auxiliary variable *z_f_* ∈ {−1, +1} that selects which of the two modes *x_f_* is in:

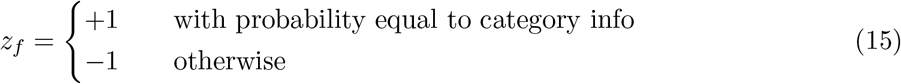

such that

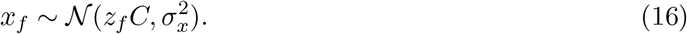

We then optimize 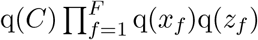.

Mean-Field Variational Bayes is a coordinate ascent algorithm on the parameters of each approximate marginal distribution. To derive the update equations for each step, we begin with the following [40]:

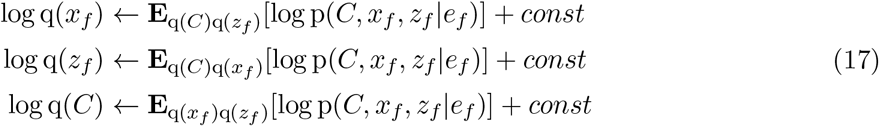

After simplifying, the new q(*x_f_*) term is a Gaussian with mean given by equation (18) and constant variance

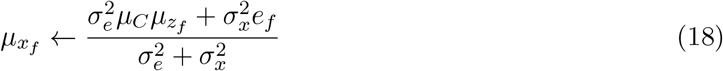

where μ_C_ and μ_z_ are the means of the current estimates of q(C) and q(z).

For the update to q(*z_f_*) in terms of log odds of *z_f_* we obtain:

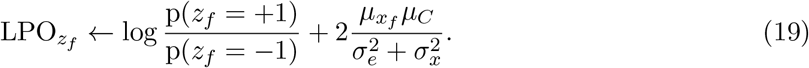

Similarly, the update to q(C) is given by:

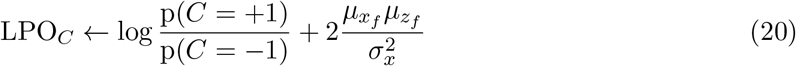

Note that the first term in equation (20) – the log prior – will be replaced with the log posterior estimate from the previous frame (see Supplemental Algorithm S2). Comparing equations (20) and (9), we see that in the variational model, the log likelihood odds estimate is given by

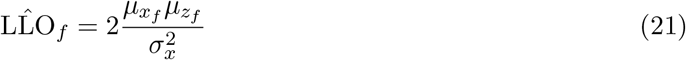

Analogously to the sampling model we assume a number of updates *n_U_* reflecting the speed of relevant computations in the brain relative to how quickly stimulus frames are presented. Unlike for the sampling model, naively amortizing the updates implied by equation (21) *n_U_* times results in a stronger primacy effect than observed in the data, since the Variational Bayes algorithm naturally has attractor dynamics built in. Allowing for an additional parameter η scaling this update (corresponding to the step size in Stochastic Variational Inference [28]) seems biologically plausible because it simply corresponds to a coupling strength in the feed-forward direction. Decreasing η both reduces the primacy effect and improves the model’s performance. Here we used η = 0.05 in all simulations based on a qualitative match with the data. The full variational model is given in Algorithm S2.

### Fitting the Extended ITB model to data

To explore alternatives, we implemented an Integration to Bound (ITB) model in our simplified 3-variable hierarchical task model, *C* → *x_f_* → *e_f_*. The dynamics of the integrator model were nearly identical to equation (11), using the exact log likelihood odds, but with added noise:

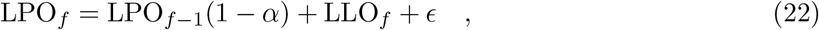

where *ϵ* is zero-mean Gaussian noise with variance 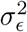 [74, 67, 6, 11, 19]. Although *α* plays a similar role to *γ* from the hierarchical inference models (both *α* and *γ* are referred to as “leak” parameters), we distinguish between them to avoid confusion. Whereas *α* < 0 produces confirmation-bias dynamics in the Extended ITB model, in the hierarchical inference models a confirmation bias occurs when *γ* is small but positive and category information is high (that is, the confirmation bias in hierarchical inference is due to biased estimation of 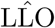 rather than to *γ*). In the Extended ITB model, whenever LPO_*f*_ crosses the bound at ±*B*, it “sticks” to that bound for the rest of the trial regardless of further evidence. Note that in the unbounded case noise does not affect the shape of the temporal weights (only their magnitude), but noise interacts with the bound to determine the shape as well as overall performance. Supplemental Figure S7 shows the performance and temporal biases of the ITB model for a range of parameter values.

Per observer per condition, we used Metropolis Hastings (MH) to infer the joint posterior over seven parameters: the category prior (*p_C_*), lapse rate (*λ*), decision temperature (*T*), integration noise (*ϵ*), bound (*B*), leak (*α*), and evidence scale (*s*). One challenge for fitting models is that the mapping from signal in the images (**S**) to “log odds” to be integrated (LLO) depends on category information, sensory information, and on unknown properties of each observer’s visual system. The evidence scale parameter, *s*, was introduced because although we can estimate the ground truth category information in each task (0.6 for HSLC and 0.9 for LSHC), the *effective* sensory information depends on each observer’s visual system and will differ between the two tasks. Using logistic regression, we explored plausible nonlinear monotonic mappings between signals **S** and log-odds, but found that none performed better than linear scaling when applied to sub-threshold trials. We therefore used LLO ≈ *g*(**S**/*s*), where *g* is a sigmoidal function that accounts for category information being less than 1, and inferred *s* jointly along with other parameters of the model. The scale s was fixed to 1 when fitting the ground-truth models, as the mapping between evidence and log odds is completely known in those cases.

Each trial, the Extended ITB model followed the noisy integration dynamics in (22), where 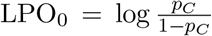 and LLO_*f*_ was computed exactly, as described above. After integration, the decision then incorporated a symmetric lapse rate and temperature:

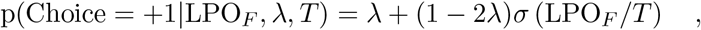

where σ(*a*) is the sigmoid function, σ(*a*) ≡ (1 + exp(−*a*))^−1^. Note that if the bound is hit, then LPO_*F*_ = ±*B*, but the temperature and lapse still apply. To compute the log likelihood for each set of parameters, we numerically marginalized over the noise, *ϵ*, by discretizing LPO into bins of width at most 0.01 between −*B* and +*B* (clipped at 3 times the largest LPO reached by the ideal observer) and computing the *probability mass* of LPO_*f*_ given LPO_*f*-1_, LLO_*f*_, and *ϵ*. This enabled exact rather than stochastic likelihood evaluations within MH.

The priors over each parameter were set as follows. p(*p_C_*) was set to Beta(2, 2). p(λ) was set to Beta(1, 10). p(*α*) was uniform in [−1, 1]. p(*s*) was set to an exponential distribution with mean 20. p(*ϵ*) was set to an exponential distribution with mean 0.25. p(*T*) was set to an exponential distribution with mean 4. p(*B*) was set to a Gamma distribution with (shape,scale) parameters (2, 3) (mean 6). MH proposal distributions were chosen to minimize the autocorrelation time when sampling each parameter in isolation.

We ran 12 MCMC chains per observer per condition. The initial point for each chain was selected as the best point among 500 quasi-random samples from the prior. Chains were run for variable durations based on available shared computing resources. Each was initially run for 4 days; all chains were then extended for each model that had not yet converged according to the Gelman-Rubin statistic, 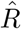 [22, 10]. We discarded burn-in samples separately per chain post-hoc, defining burn-in as the time until the first sample surpassed the median posterior probability for that chain (maximum 20%, median 0.46%, minimum 0.1% of the chain length for all chains). After discarding burn-in, all chains had a minimum of 81k, median 334k, and maximum 999k samples. Standard practice suggests that 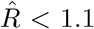 indicates good enough convergence. The slowest-mixing parameter was the signal scale (*s*), with 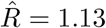 in the worst case. All 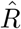 values for the parameters relevant to the main analysis − *α*, *B*, and *β* – indicated convergence ([min, median, max] values of 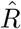 equal to [1, 1.00335, 1.032] for *α*, [1.0005, 1.00555, 1.0425] for *B*, and [1, 1.0014, 1.0178] for all *β* values in ablation analyses.

### Estimating temporal slopes and ablation indices implied by model samples

To estimate the the shape of temporal weights implied by the model fits, we simulated choices from the model once for each posterior sample after thinning to 500 samples per chain for a total of 6k samples per observer and condition. We then fit the slope of the exponential weight function, *β*, to these simulated choices using logistic regression constrained to be an exponential function of time as described earlier (equation (3)). This is the *β*_fit_ plotted on the y-axis of Figure 6b. For the ablation analyses, we again fit *β* to choices simulated once per posterior sample of model parameters, but setting *α* = 0 in one case or (*B* = ∞, *ϵ* = 0) in the other.

We used a hierarchical regression analysis to compute “ablation indices” per observer and per parameter. The motivation for this analysis is that observers have different magnitudes of primacy and recency effects, but the *relative* impact of the leak or bound and noise parameters appeared fairly consistent throughout the population (Supplemental Figure S12), so a good summary index measures the *fraction* of the bias attributable to each parameter, which directly relates to the slope of a regression line through the origin. To quantify the net effect of each ablated parameter per observer, we regressed a linear model with zero intercept to *β*_fit_ versus *β*_true_. If an ablated parameter has little impact on *β*_fit_, then the slope of the regression will be near 1, so we use 1 minus the linear model’s slope as an index of the parameter’s contribution. The regression model accounted for errors in both *x* and *y* but approximated them as Gaussian. Defining *m* to be the regression slope for the population and *m_i_* to be the slope for observer *i*, the regression model was defined as

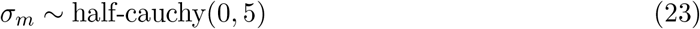

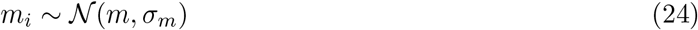

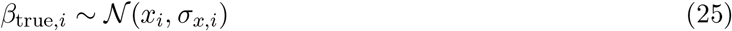

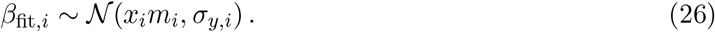

This model was implemented in STAN and fit using NUTS [14]. The regression was done separately for each experimental condition and each set of ablated parameters. Equations (23) and (24) are standard practice in hierarchical regression – they capture the idea that there is variation in the parameter of interest (the slope *m*) across observers which is normally distributed with unknown variance, σ_*m*_, but that this variance is encouraged to be small if supported by the data. The variable *x_i_* is the “true” x location associated with each observer, which is inferred as a latent variable to account for measurement error in both x (25) and y (26) dimensions. Measurement errors in *β*_true_, σ*_x,i_* were set to the standard deviation in *β* across bootstraps. Measurement errors in *β*_fit_, σ_*y,i*_ were set to the standard deviation of the posterior predictive distribution over *β* from simulated choices on each sample of model parameters as described above. We set *x_i_* and *y_i_* to the median values of *β*_true_ (across bootstrapped trials) and *β*_fit_ (across posterior samples), respectively.

### Ground-truth models

Based on observations of the temporal weighting profile alone, the transition between primacy and recency could be explained by bounded integration with a changing leak amount in the LSHC condition and high bound in the HSLC condition (Supplemental Figure S7g-i). To verify that all of the above fitting and ablation procedures could distinguish a confirmation bias from bounded integration, we tested them on two ground-truth models: one where choices were simulated from a hierarchical inference (IS) model, and one where choices were simulated from an ITB model. All ground-truth parameter values are given in Supplemental Table S3, which were chosen to meet two criteria: first, constant performance at 70% in both LSHC and HSLC regimes, and second, matched temporal slopes (a primacy effect with shape *β* ≈ −0.1 in the LSHC condition and a recency effect with shape *β* ≈ 0.1 in the HSLC condition for both models). This analysis confirmed that bounded integration is indeed distinguishable from a confirmation bias (*α* < 0), in terms of the quality of the fit (Supplemental Figures S9), different inferred parameter values (Supplemental Figure S10), and the ablation tests (Supplemental Figure S11).

## Acknowledgements

This work was supported by NEI/NIH awards R01 EY028811-01 (RMH) and T32 EY007125 (RDL, JLY), as well as an NSF/NRT graduate training grant NSF-1449828 (RDL).

## Author Contributions

Author contributions are shown in the following table, where black = significant contribution, gray = partial contribution, and white = zero or minimal contribution.

|  | RL | AC | JB | JY | RH |
| --- | --- | --- | --- | --- | --- |
| Experiment Design | black | black | white | black | black |
| Experiment Code | black | gray | white | white | white |
| Data Collection | gray | black | white | white | white |
| Data Analysis | black | gray | white | gray | gray |
| Sampling Model | black | black | white | white | black |
| Variational Model | black | white | black | white | white |
| ITB Model + fitting | black | white | white | white | gray |
| Writing | black | gray | gray | gray | black |

