## Supplemental text and figures for "A confirmation bias in perceptual decision-making due to hierarchical approximate inference"

#### Sensory Information and Category Information in Previous Literature

In this section we justify our categorization of previous studies' stimuli into the low-sensory/high-category information (LSHC) or high-sensory/low-category information (HSLC) regime in relation to Figure 1c and Table S1.

##### Studies finding a primacy effect

Kiani et al. (2008) studied the classic motion direction discrimination task in which a monkey views a dynamic random dot motion stimulus with a certain percentage of "coherent" dots moving together and the rest moving randomly [16, 18]. Monkeys were trained to categorize the direction of motion as predominantly leftward or rightward. Since the direction of the coherently moving dots (the signal) does not change over time within a trial, this stimulus contains high category information. Since the motion direction is difficult to perceive for any motion frame, it contains low sensory information [16].

Nienborg et al. (2009) developed a task in which observers viewed a disc with varying binocular disparity. The disc moved back and forth relative to a reference plane (the surrounding ring), changing every 10ms, at a rate too high for the macaques' (and humans') binocular system to resolve, resulting in a percept of a jittering cloud of dots which was located slightly in front of or behind the surrounding ring and blurred in depth (Nienborg – private communication). After 200 frames presented over 2 seconds, observers judged whether the center disc was in front or behind the reference plane. Since the location of the perceived dot cloud is relatively stable, but itself uncertain with respect to the reference, this stimulus contains high category and low sensory information [19].

##### Studies finding a recency effect or flat weighting

In two similar studies by Wyart et al. (2012) and by Drugowitsch et al. (2016), human participants viewed a sequence of eight clearly visible oriented gratings presented for at least 250ms each. Participants reported whether, on average, the tilt of the eight elements fell closer to the cardinal or diagonal axes. These tasks contain high sensory information since for a observer there is little uncertainty about the orientation of any one grating. However they contain low category information since the orientation of any one grating provides only little information about the correct choice [25, 11].

Brunton et al. (2013) studied both a visual task and an auditory task where observers were trained to indicate whether they saw/heard more flashes/clicks on the left or right side of the midline. These task stimuli contain high sensory information since each flash/click is high contrast/loud – well above observers' detection thresholds. However, they contain low category information since each flash/click contains only little information about the correct choice [8].

#### Stimulus details

The stimulus was constructed from white noise that was then masked by a kernel in the Fourier domain to include energy at a range of orientations and spatial frequencies but random phases [3, 20, 6]. The Fourier-domain kernel consisted of a product of two probability density functions (PDFs): a von Mises PDF over orientation, and a Rician PDF over spatial frequency. This is best expressed using polar coordinates in the Fourier domain:

$$K_{\rho\theta} = \text{vonMises}(\theta; \mu_\theta, \kappa)\text{Rician}(\rho; \mu_\rho, \sigma_\rho)$$

where $\theta$ is the angular coordinate and $\rho$ is the spatial frequency coordinate. After transforming back from the Fourier domain to an image, we applied a soft circular aperture with a hole cut out in the center for the fixation cross. The full pixel-space mask is defined by the equation

$$M = \underbrace{\exp(-4\hat{\rho}^2)}_{\text{Gaussian aperture}} \times \underbrace{(1 + \text{erf}(10 \times (\hat{\rho} - \tau_{\text{ap}}/w_{\text{im}})))}_{\text{Center cutout for fixation cross}}$$

where $\hat{\rho}$ is the normalized Euclidean distance to the center of the image ($\hat{\rho} = 0$ at the center, and $\hat{\rho} = \sqrt{2}$ at the corners), and erf is the Error Function. $\tau_{\text{ap}}$ controlled the width of the central cutout, and $w_{\text{im}}$ is the total width of the stimulus. To summarize, each stimulus frame, I, was generated according to

$$\text{I} = M \otimes \mathcal{F}^{-1}\left[\mathcal{F}[\mathcal{W}] \otimes K_{\rho\theta}\right]$$

where $\mathcal{F}$ is the 2D discrete Fourier transform, $\otimes$ is element-wise multiplication of each pixel, and $\mathcal{W}$ is white noise. Images were displayed using Psychtoolbox on a 1920x1080px 120

Hz monitor with gamma-corrected luminance [7]. Using an 8-bit luminance range (0 to 255), each frame was normalized to $127 \pm c$ where $c$ is a contrast parameter. All stimulus parameters are summarized in table S2.

### Algorithms

**Algorithm S1** Importance Sampling (IS) model for evidence integration

$\text{LPO} \leftarrow \log \frac{\text{p}(C=+1)}{\text{p}(C=-1)}$ ▷ initialize log posterior odds to log prior odds
**for** $f = 1$ to $F$ **do**
  **for** $n = 1$ to $n_\text{U}$ **do**
    $p_C \leftarrow (1 + \exp(-LPO))^{-1}$ ▷ current posterior that $C = +1$
    $\hat{p}(x) \leftarrow p_C \mathcal{N}(+1, \sigma_x^2) + (1 - p_C)\mathcal{N}(-1, \sigma_x^2)$ ▷ Mixture of Gaussians prior on $x$
    $Q(x) \leftarrow \hat{p}(x)p(e_f|x)$
    **for** $s = 1 \ldots S$ **do**
      $x^{(s)} \sim Q(x)$ ▷ sensory sample from current posterior
      $p_+^{(s)} \leftarrow p(x^{(s)}|C = +1)$ ▷ contribution of each sample to $C = +1$ pool
      $p_-^{(s)} \leftarrow p(x^{(s)}|C = -1)$ ▷ contribution of each sample to $C = -1$ pool
      $w^{(s)} \leftarrow \left(\sum_c \text{p}(x^{(s)}|C = c)\text{p}_{f-1}(C = c)\right)^{-1}$ ▷ (unnormalized) weight of each sample
    **end for**
    $w \leftarrow w / \sum_{s'} w^{(s')}$ ▷ (optionally) normalize weights
    $p_+^{tot} \leftarrow \sum_s p_+^{(s)} w^{(s)}$ ▷ aggregate evidence for $C = +1$
    $p_-^{tot} \leftarrow \sum_s p_-^{(s)} w^{(s)}$ ▷ aggregate evidence for $C = -1$
    $\hat{\text{LLO}}_f \leftarrow \log p_+^{tot} - \log p_-^{tot}$
    $\text{LPO} \leftarrow \text{LPO}(1 - \gamma/n_\text{U}) + \hat{\text{LLO}}_f/n_\text{U}$ ▷ equations (14,11) amortized for $n_\text{U}$ updates
  **end for**
**end for**

**Algorithm S2** Variational Bayes (VB) model for evidence integration

LPO $\leftarrow \log \frac{p(C=+1)}{p(C=-1)}$ ▷ initialize to log prior odds
**for** $f = 1$ to $F$ **do**
  $\mu_{z_f} \leftarrow 2p(z_f = +1) - 1$ ▷ initialize $\mu_{z_f}$ to the prior
  **for** $n = 1$ to $n_U$ **do**
    $\mu_C \leftarrow 2(1 + \exp(-\text{LPO}_C))^{-1} - 1$ ▷ convert log-odds to mean of $C$
    $\mu_{x_f} \leftarrow \frac{\sigma_e^2 \mu_C \mu_{z_f} + \sigma_x^2 e_f}{\sigma_e^2 + \sigma_x^2}$ ▷ equation (18)
    $\text{LPO}_{z_f} \leftarrow \log \frac{p(z_f=+1)}{p(z_f=-1)} + 2\frac{\mu_{x_f}\mu_C}{\sigma_x^2+\sigma_e^2}$ ▷ equation (19)
    $\mu_{z_f} \leftarrow 2(1 + \exp(-\text{LPO}_{z_f})^{-1} - 1$ ▷ convert log-odds to mean of $z_f$
    $\hat{\text{LLO}}_f \leftarrow \frac{2\mu_{x_f}\mu_{z_f}}{\sigma_x^2}$ ▷ Equation (21)
    LPO $\leftarrow$ LPO$(1 - \gamma/n_U) + \eta \hat{\text{LLO}}_f / n_U$ ▷ Equations (11) and (20) amortized for $n_U$ updates with update strength $\eta$
  **end for**
**end for**

### Optimal bias correction

A leak term approximates optimal inference in a changing environment when total evidence is weak [13], but each trial of our task is stationary. One might therefore expect that a leak term, or $\gamma > 0$, would impair the model's performance in our task. On the other hand, we motivated the leak term by suggesting that it could approximately correct for the confirmation bias. Under this second interpretation, one might instead expect performance to *improve* for some $\gamma > 0$, especially for conditions where the confirmation bias was strong.

We investigated the relationship between the leak ($\gamma$) and model performance. First, we simulated the importance sampling model with $\gamma = 0.1$ and $\gamma = 0.5$ and compared its performance across the space of category and sensory information (Figure S5a-b). We found that in the LSHC regime where the confirmation bias had been strongest, the larger value of $\gamma$ counteracts the bias and leads to better performance, but in the HSLC regime where there had been no confirmation bias, the optimal $\gamma$ is zero (Figure S5c). We thus see that the optimal value of $\gamma$ depends on the task statistics, i.e. the balance of sensory information and category information: the stronger the primacy effect or confirmation bias, the higher $\gamma$ must be to correct for it (Figure S5d). Analogous results were found for the variational model (Figure S6).

We next asked what the effect would be on the model's temporal weights if it could utilize the best $\gamma$ for each task. We found that the $\gamma-$optimized model displayed near-flat weights across the entire space of tasks (Figure S5f). Our experimental data therefore imply that either the brain does not optimize its leak to the statistics of the current task, or that it does so on a timescale that is slower than a single experimental session (roughly 1hr, Methods).

### Detailed comparison with integration to bound (ITB)

The primary alternative explanation for primacy effects in fixed-duration integration tasks proposes that observers integrate evidence to an internal *bound*, at which point they cease paying attention to the stimulus [16]. Because the bound is crossed at different times on different trials, the average weight observers give to each frame is a decreasing function of the frame number, i.e. a primacy effect. We implemented an integration-to-bound (ITB) observer in our hierarchical inference framework and replicated the observation that bounded and noisy integration results in primacy effects (Figure S7a-b, using $\sigma_x^2 = 0.1$, $\epsilon = 0.35$, $\alpha = 0$, and $B = 1.2$). Importantly, this mechanism depends only on the net log likelihood per frame regardless of how it is partitioned into category information and sensory information. Classic ITB therefore always predicts the same temporal weights as long as performance is held constant. ITB does, however, predict a change in temporal weighting as a function of task difficulty, because the bound is hit earlier in a trial when evidence is stronger (Figure S7c). However, this explanation is unlikely to explain the changes seen on our data given that our experiment used a continuous staircase procedure which sustained performance near 70% in both tasks.

We next investigated the behavior of a *leaky*, noisy, and bounded integrator. While the addition of a leak term shifts the effective weights in the direction of a recency effect, we again see no systematic changes across the space of tasks (Figure S7d-f). In order to produce different regimes of temporal biases at fixed performance levels, then, either the bound, the leak term, or both must change as a function of category information and sensory information. We next simulated a leaky ITB model in which the leak term, $\alpha$, varied with category information: small but positive $\alpha$ in the LSHC regime and large $\alpha$ in the HSLC regime. This change is plausible because observers may adopt a strategy that discounts past evidence more when the world appears more volatile [13]. This model is dominated by bounded integration in the LSHC condition and by leaky integration in the HSLC condition, qualitatively reproducing the trends in our data (Figure S7g-i).

There are thus two families of models in qualitative agreement with our observers' data: hierarchical inference with a confirmation bias, or bounded integration with a leak that depends on the task. Both model families explain recency effects as the result of leaky integration but differ in their account of primacy effects. We reasoned that these models might be distinguished using data from our LSHC condition: whereas they agree on the sign and magnitude of the temporal bias as measured by an exponential fit $\beta$, they make divergent predictions for observers' confidence, determined by the magnitude of the integrated log odds at the end of a trial. According to the confirmation-bias mechanism, observers should count all evidence in a trial but *over*-count early evidence, inflating their confidence relative to an unbiased integrator. According to the ITB mechanism, however, the magnitude of the bound itself sets an upper limit on log odds, and thus an upper limit on confidence, truncating the range of confidences relative to an unbiased integrator. Because we did not ask observers to report confidence in their choices, these predictions cannot be tested directly. However, this line of reasoning suggests that these mechanisms may nonetheless be distinguished by fitting models to observers' data; confident choices are predictable choices.

We first tested whether the two primacy mechanisms – a confirmation bias or bounded integration – are quantitatively distinguishable in ground-truth data. We simulated choices from the ground-truth IS and ITB models already described (parameters given in Table S3; respective models plotted in Figure 3c-e and Figure S7g-i). The models were matched both in performance and in their temporal biases, exhibiting a primacy effect ($\beta \approx -0.1$) in the LSHC condition and a recency effect ($\beta \approx +0.1$) in the HSLC condition. Due to the internal stochasticity of the IS model, it is infeasible to infer its parameters directly. However, we found that an ITB model with a large bound and negative leak ($\alpha < 0$) is *functionally* indistinguishable from the IS model (Figure S9). Recall that a leak term, $\gamma$, was introduced in equation (11) and explains recency effects when $\gamma > 0$. The analogous term in the ITB model is $\alpha$, which similarly produces recency effects when it is positive. When $\alpha < 0$, however, this has the opposite effect of amplifying already accumulated evidence, leading to a primacy effect due to a mechanism that is *functionally* equivalent to a confirmation bias [9, 5]. The key question thus becomes: are the primacy effects in our data better explained by a negative leak term or by bounded integration [23]? These mechanisms not mutually exclusive and in principle both may contribute. We therefore fit a single ITB model with $-1 < \alpha < 1$ to each condition. By fitting a single model that contains both mechanisms as special cases, we compare them on equal terms. In order to estimate the relative contribution of each mechanism, we used MCMC sampling to infer the full posterior over all parameters.

We verified that these two distinct parameter regimes – negative leak or bounded integration – are distinguishable in ground-truth data. Indeed, in the case of the IS model, the posterior concentrated on unbounded integration with $\alpha < 0$ in the LSHC condition and unbounded but leaky integration in the HSLC condition. In the case of ground truth data from the ITB model in Figure S7g-i, the posterior concentrated around the ground truth parameters (Figure S10). We further verified that ablating either the leak ($\alpha$), or ablating bound and noise parameters together, led to distinguishable patterns of temporal biases in the ground-truth models (Figure S11).

### Simulation of a larger hierarchical inference model

We simulated the hierarchical sampling-based inference model of [15]. Unlike our reduced $C \to x \to e$ models in the main text with only scalar variables, the model of [15] decomposes as $C \to \mathbf{G} \to \mathbf{X} \to \mathbf{I}$ where $\mathbf{I}$ is an entire image, and $\mathbf{X}$ and $\mathbf{G}$ represent entire populations of V1 and V2 neurons respectively. We will refer to this as the HBF16 model in what follows. Trying to better understand inference dynamics and the source of primacy effects in the HBF16 inspired the present work. In particular, the original model was shown to produce primacy effects in a task which we would now categorize as having low-sensory and high-category information.

The original HBF16 model was run on a coarse orientation discrimination task between low-contrast vertical and horizontal gratings embedded in white noise with variance 1. As in our reduced models in the main text, we adapted the generative model to the statistics of the stimuli as we transitioned from LSHC to HSLC conditions. In the main text, we converted sensory information into the variance of two Gaussians centered at $\pm 1$. In the HBF16 model, sensory information is instead determined by the *contrast* of a stimulus with fixed noise. We therefore made no change to the *generative* structure of $\mathbf{X} \to \mathbf{I}$ because higher contrast images immediately results in higher signal to noise in $\mathbf{X}$. We manipulated category information in the stimulus, as in the models in the main text, by randomly flipping the orientation of each of the 10 frames per trial with probability $p_{match}$. Lower category information in the stimulus requires a weaker coupling from $\mathbf{G}$ to $C$, parameterized by $\kappa$. For each V2-like grating element $G_i$ with preferred orientation $\theta_i$, the generative model couples $C$ to $\mathbf{G}$ as follows:

$$\mathrm{p}(G_i = 1|C) \propto \begin{cases} \exp(\kappa \cos(\theta_i - \theta_{C=1})) & \text{if } C = 1 \\ \exp(\kappa \cos(\theta_i - \theta_{C=2})) & \text{if } C = 2 \end{cases} \tag{1}$$

where $\theta_{C=c}$ is the true grating orientation for category $c \in \{1, 2\}$. Note that each $G_i$ is binary, indicating the presence or absence of a grating element (see [15] for additional details). Clearly, as $\kappa$ goes to zero, $C$ and $\mathbf{G}$ become independent, and as $\kappa$ gets large, $C$ uniquely determines which grating orientation is present, and, conversely, samples of $\mathbf{G}$ strongly determine $C$. Thus $\kappa$ controls the strength of the positive feedback or confirmation bias in this model.

The strength of the coupling between $C$ and $\mathbf{G}$ is naturally quantified with the ROC of the two cases of von Mises distributions in (1). As in the main paper, this quantifies category information (in the generative model rather than in the stimulus) on a scale between 0.5 and 1. Denoting this function as $\mathrm{p} = \mathrm{roc}(\kappa)$ and its inverse as $\kappa = \mathrm{roc}^{-1}(\mathrm{p})$, we set $\kappa$ in our simulations to $\mathrm{roc}^{-1}(\mathrm{p}_{match})/\mathrm{roc}^{-1}(0.9)$. This way, $\kappa$ scaled appropriately with the amount of information in the stimulus, and $\kappa = 1$ when category information is 0.9 to approximately the original parameter regimes of HBF16.

We additionally extended the model of HBF16 to include a leak parameter in the update to the log odds of $C$, and set the leak to 0.01 in the simulations (equivalent to $\gamma = 0.08$ in the main paper where we divided $\gamma$ by the number of updates per frame). We simulated 200 trials from the HBF16 model across a range of contrast values from 0 to 10 and $\mathrm{p}_{match}$ values ranging from 0.51 to 0.99. We then smoothed the resulting performance grid and plotted the results in Figure S8a, and recapitulates the patterns seen in our reduced models. We selected two points in this space – corresponding to one LSHC and one HSLC condition – for 5000 additional trials. We then computed temporal weights using AR2-regularized logistic regression. Results are plotted in Figure S8b, showing a transition from primacy in the LSHC condition to recency in the HSLC condition. (Note that without any leak, the HBF 16 model only transitions to flat weights in the HSLC condition but requires higher sensory information for equivalent LSHC performance, exactly as in our reduced models; not plotted). This demonstrates that our insights from the reduced hierarchical inference models used in the main text can generalize to larger hierarchical inference settings with a large number of variables and nontrivial dynamics.

### Additional model-fitting details

To determine whether observers' strategies were better described by confirmation bias dynamics or bounded integration, we initially sought to use standard *model comparison* methods. Ideally, Bayesian model comparison is done by computing Bayes Factors, or the ratio of the marginal likelihoods of the data under two models being compared [4]. The marginal likelihood may be estimated by procedures similar to cross-validation [12], which requires repeatedly performing full Bayesian inference over model parameters conditioned on random splits or subsets of the full dataset. For this to be feasible, the "inner loop" of Bayesian inference must be efficient. The primary barrier to this approach is the fact that the likelihood in the IS model is only known implicitly through stochastic simulations. Simulation-based inference methods are an active area of research [24, 14, 17, 21, 22, 1].

For all of our models, the likelihood of the observer's choice on trial $t$, written $\mathrm{p}(\mathrm{choice}_t|\mathbf{S}_t, \theta)$ for stimulus sequence $\mathbf{S}_t$ and model parameters $\theta$, is the Bernoulli probability of the observed choice given the model's confidence on the final frame, *marginalizing over the internal stochasticity of the model.* That is, for a fixed stimulus $\mathbf{S}_t$ and parameters $\theta$, the model may output a different final log odds, $\mathrm{LPO}_F$, on multiple runs. The likelihood can be written

$$\mathrm{p}(\mathrm{choice}_t|\mathbf{S}_t, \theta) = \int_{-\infty}^{\infty} \underbrace{\mathrm{p}(\mathrm{choice}_t|\mathrm{LPO}_F, \theta)}_{(i)} \underbrace{\mathrm{p}(\mathrm{LPO}_F|\mathbf{S}_t, \theta)}_{(ii)} \mathrm{dLPO}_F \quad .$$

The first term, $(i)$, is the lapse- and temperature-adjusted probability of making a choice given a final confidence or belief value of $\mathrm{LPO}_F$. The second term, $(ii)$, depends on the internal stochasticity of the model. In the case of ITB models, all internal stochasticity is due to the integration noise $\epsilon$, and can be numerically marginalized by internally maintaining a *distribution* of possible log posterior odds each frame, and updating that distribution for each frame, computing a new distribution, $\mathrm{p}(\mathrm{LPO}_f|\mathrm{LPO}_{f-1}, \mathrm{LLO}_f, \theta)$, taking into account the total probability mass that has crossed the bounds $\pm B$. This is precisely how we estimate the likelihood for the Metropolis Hastings sampler used in the main text. We cannot, however, apply the same trick to the IS model. Whereas the ITB models' internal stochasticity is simply additive Gaussian noise with variance $\epsilon^2$, internal stochasticity in the IS model comes from the location of generated samples in the SNIS algorithm. If drawing $S = 5$ samples per update, as in our main simulations, then marginalization would require integrating over $\mathbb{R}^5$. In general, the marginalization problem grows exponentially with $S$, which is a parameter we would in principle like to infer and may be large. As a final comment before discussing alternatives, we note that SNIS with $S$ samples can be viewed as implicitly defining a 1-dimensional distribution over $x$ after $S-1$ marginalization steps [10]; however, this distribution is not known in closed form (or whether it has a closed form), and we were unable to derive a sub-exponential-time expression for numerically approximating it.

An cheaper alternative approach to model comparison, compared to performing full Bayesian inference in an inner-loop, is to search for the maximum likelihood or maximum a posteriori estimate of the parameters (MLE or MAP), then approximately correct for biases by adjusting the model score by the number of parameters (as in AIC) or the number of parameters and amount of data (as in BIC). *Search* methods with a stochastic objective are in general more mature than *inference* methods with stochastic likelihood evaluations, suggesting this may be a promising approach. It requires two ingredients: a method to get unbiased (but possibly variable) estimates of the log likelihood, and a method to search for the maximum of a noisy objective. We implemented the Inverse Binomial Sampling (IBS) method of van Opheusden et al (2020) to get unbiased but noisy log likelihood estimates . Briefly, IBS estimates the likelihood of each trial by counting the number of repeated (stochastic) simulations it takes before the model makes the same choice as the observer. Let $k_t$ be the number of simulations before the first match, then IBS estimates the log likelihood for that trial as

$$\hat{LL}_t = \psi(1) - \psi(k_t) \quad , \tag{2}$$

where $\psi$ is the digamma function [24]. Crucially, $\hat{LL}_t$ is an *unbiased* estimator of the true $LL_t$. Other naive methods derived by considering how to estimate the likelihood directly (as opposed to the log likelihood) result in biases after taking the log. The full log-likelihood estimate is given by $\hat{LL} = \sum_{t=1}^{T} \hat{LL}_t$. Its variance grows with the number of trials, so we averaged together $\sqrt{T}$ repeats of the IBS estimator per evaluation. With an unbiased estimator of the log likelihood in hand, we used Bayesian Adaptive Direct Search (BADS) to search for the maximum likelihood parameters [2]. We began with a quasi-random grid of 5k points sampled from the prior over each parameter and evaluated their estimated log likelihood. For each BADS run, we perturbed the set of evaluated log likelihoods by adding Gaussian noise proportional to the empirical standard deviation of $\hat{LL}$ (i.e. Thompson Sampling), then selected the maximum as the starting point. We re-ran this procedure for at least 20 and at most 1000 searches (stopping when enough runs agreed on the value of $\hat{LL}$ at the MLE). Using the best estimate of $\hat{LL}$ for each model and condition, re-estimated with $10\sqrt{T}$ repeats of IBS, we computed AIC:

$$\text{AIC} = -2\hat{LL} + 2P \quad ,$$

where $P$ is the number of parameters in the model. Because $\hat{LL}$ is stochastic with known empirical variance, we plotted AIC for each model fit to ground-truth data with error bars in Figure S9.

Ultimately, our conclusions from this AIC-based comparison on ground-truth models were two-fold. First, although we were able to recover the ground-truth parameters in each case, this method gives no sense of the *uncertainty* over those parameters, which is crucial for answering the question posed in the main text of the *extent* to which either of two mechanism produces primacy effects. Second, we observed that although the standard ITB model is distinguishable from the IS model with the constraint of a positive leak ($0 < \alpha < 1$) enforced, allowing negative leak ($-1 < \alpha < 1$) it is no longer distinguishable (Figure S9). In other words, this means that a negative leak is *functionally* indistinguishable from the IS model in the LSHC condition. Further, the same ITB model family with a positive leak is *functionally* indistinguishable from the IS model in the HSLC condition. Taken together, this implies that the key question of whether primacy effects are due to bounded integration or due to self-reinforcing dynamics when integrating LPO can be answered even more directly and more fairly by comparing *parameter regimes* within the ITB model family with negative leak rather than comparing across model families of IS and ITB. For this reason, we pursued full inference over ITB model parameters in the main text rather than fitting a point estimate of the IS model directly to data.

| Example study | Justification for placement in task space (Figure 1, color-coded) | Suggested stimulus manipulation to change weighting (color-coded) |
|---|---|---|
| Brunton et al. (2013), Raposo et al. (2014) | Each click is perceptually clear but only weakly predictive of which side has the higher rate. | Make clicks softer or embed them in noise and increase difference in rates between left and right side. |
| Wyart et al. (2012), Drugowitsch et al. (2016) | Orientation of each frame is clear but only weakly predictive of which "deck" the orientations were drawn from. | Decrease contrast of each frame or increase pixel noise and reduce variance of orientations within each deck. |
| Kiani et al. (2008) | Net motion is weak (low coherence) and constant over a trial. | Increase motion coherence but vary net motion direction across stimulus frames within a trial. |
| Nienborg et al. (2009) | Percept is of a jittering cloud of dots whose depth is close to fixation point. | Increase the distance between cloud and fixation point in depth; vary distance across stimulus frames at a rate resolvable by depth perception |

Table S1: Justification of placement of example prior studies in Figure 1c and description of stimulus manipulations that will move it to the opposite side of the category–sensory–information space. Each manipulation corresponds to a prediction about how temporal weighting of evidence should change from primacy (red) to flat/recency (blue), or vice versa, as a result.

| **Parameter** | **Description** | **Values (Units)** |
|---|---|---|
| $\mu_\rho$ | mean spatial frequency | 6.90 (cycles per degree) |
| $\sigma_\rho$ | spread of spatial frequency | 3.45 (cycles per degree) |
| $\kappa$ | (inverse) spread of orientation energy | $0 \leq \kappa \leq 0.8$ |
| c | image contrast | 22 |
| $\tau_{\text{ap}}$ | width of central annulus cutout | 25 (pixels) or 0.43 (°) |
| $w_{\text{im}}$ | full image width & height | 120 (pixels) or 2.08 (°) |

Table S2: Stimulus parameters.

| | LSHC | | | | | | | | HSLC | | | | | | | |
|---|---|---|---|---|---|---|---|---|---|---|---|---|---|---|---|---|
| Model | SI | CI | S | $\gamma/\alpha$ | B | $\epsilon$ | T | $\lambda$ | SI | CI | S | $\gamma/\alpha$ | B | $\epsilon$ | T | $\lambda$ |
| IS | .65 | .91 | 5 | .1 | $\infty$ | 0 | .1 | 0 | .91 | .63 | 5 | .1 | $\infty$ | 0 | .1 | 0 |
| ITB | .65 | .91 | | .09 | 1.2 | .35 | .1 | 0 | .91 | .65 | | .35 | 1.2 | .35 | .1 | 0 |

Table S3: Parameters of ground-truth models used to test model-fitting. SI = sensory information. CI = category information. $\gamma/\alpha$ = leak. S = samples per batch (IS model only). B = bound (ITB model only). $\epsilon$ = integration noise. T = decision temperature. $\lambda$ = lapse rate.

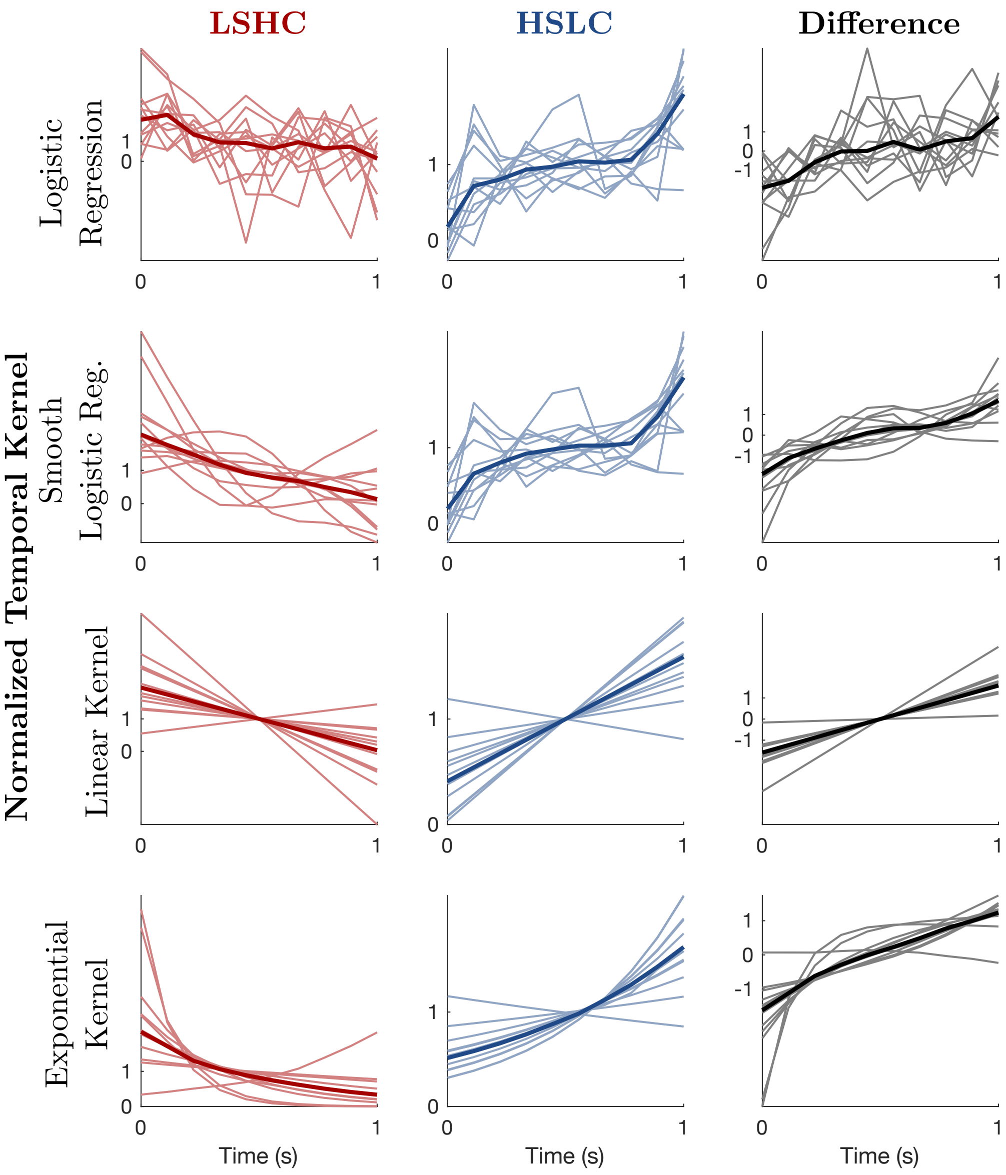


Figure S1: Temporal kernels for each condition (LSHC and HSLC), and their difference between conditions, for each of four regularization techniques. In all panels, weights are normalized to have a mean of 1, individual observers are shown as faint thin lines, and the average across observers as a darker bold line. First row ("Logisitic Regression") is the result of ridge regression for predicting choices from per-frame signal levels with no further regularization. Second row ("Smooth Logistic Regression") includes a second-order autoregressive penalty, resulting in smoother kernels. Third row ("Linear Kernels") is a three-parameter model that constrains weights to be a linear function of time. The three parameters control the slope and intercept of the kernel, and the choice bias (Methods). Fourth row ("Exponential Kernels") is a similar three-parameter model that instead constrains weights to be an exponential function of time.

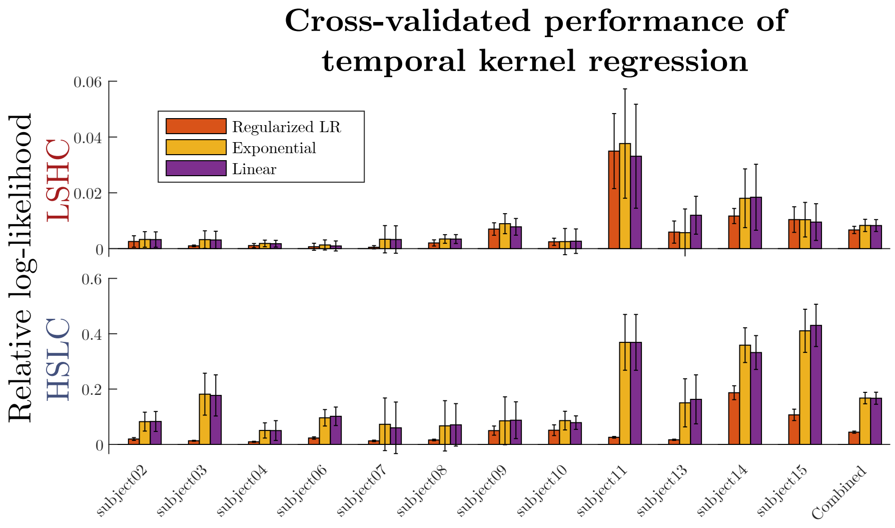


Figure S2: Cross-validation selects linear or exponential shapes for temporal weights, compared to both unregularized and smoothness-regularized logistic regression. Panels show 20-fold cross-validation performance of four regression methods to predict choices from subthreshold trials, separated by task type and by observer. All values are relative to the log-likelihood, per fold, of the unregularized model. Error bars show standard error of the mean difference in performance across folds of shuffled data. "Unregularized LR" refers to standard ridge regression with no regularization of the temporal shape. "Regularized LR" refers to the AR2-penalized logistic regression objective, where the hyperparameters were chosen to maximize cross-validated fitting performance separately for each observer. "Exponential" is the 3-parameter model where weights are an exponential function of time (equation (3) plus a bias term). Similarly, the "Linear" model constrains the weights to be a linear function of time as in equation (4), plus a bias term.

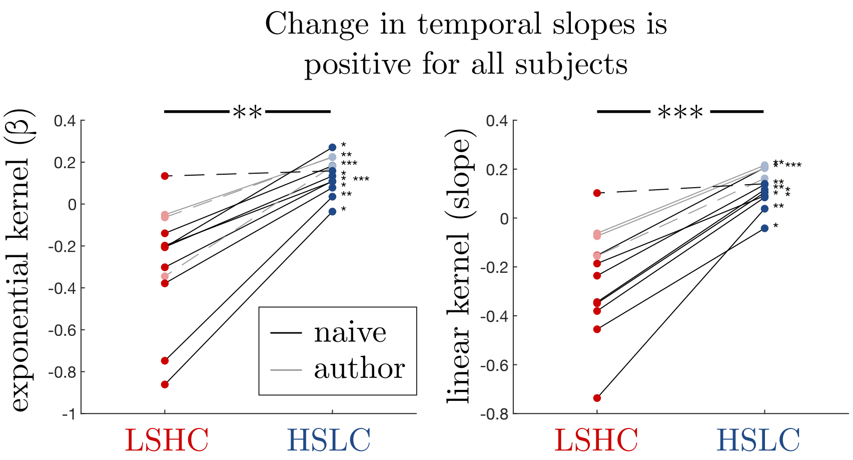


Figure S3: Left panel is the same as Figure 5e in the main text, comparing slope of temporal weights by constraining weights to be an exponential function of time. The right panel shows the same analysis with weights constrained to be a linear function of time. In both cases, 9 of 12 observers individually have a significant increase in slope ($p < 0.05$, bootstrap). A one-sided sign test on the medians for each observer reveals a significant population effect with $p = .0032$ ($**$)for the exponential method and $p = 0.00024$ ($***$) for the linear method.

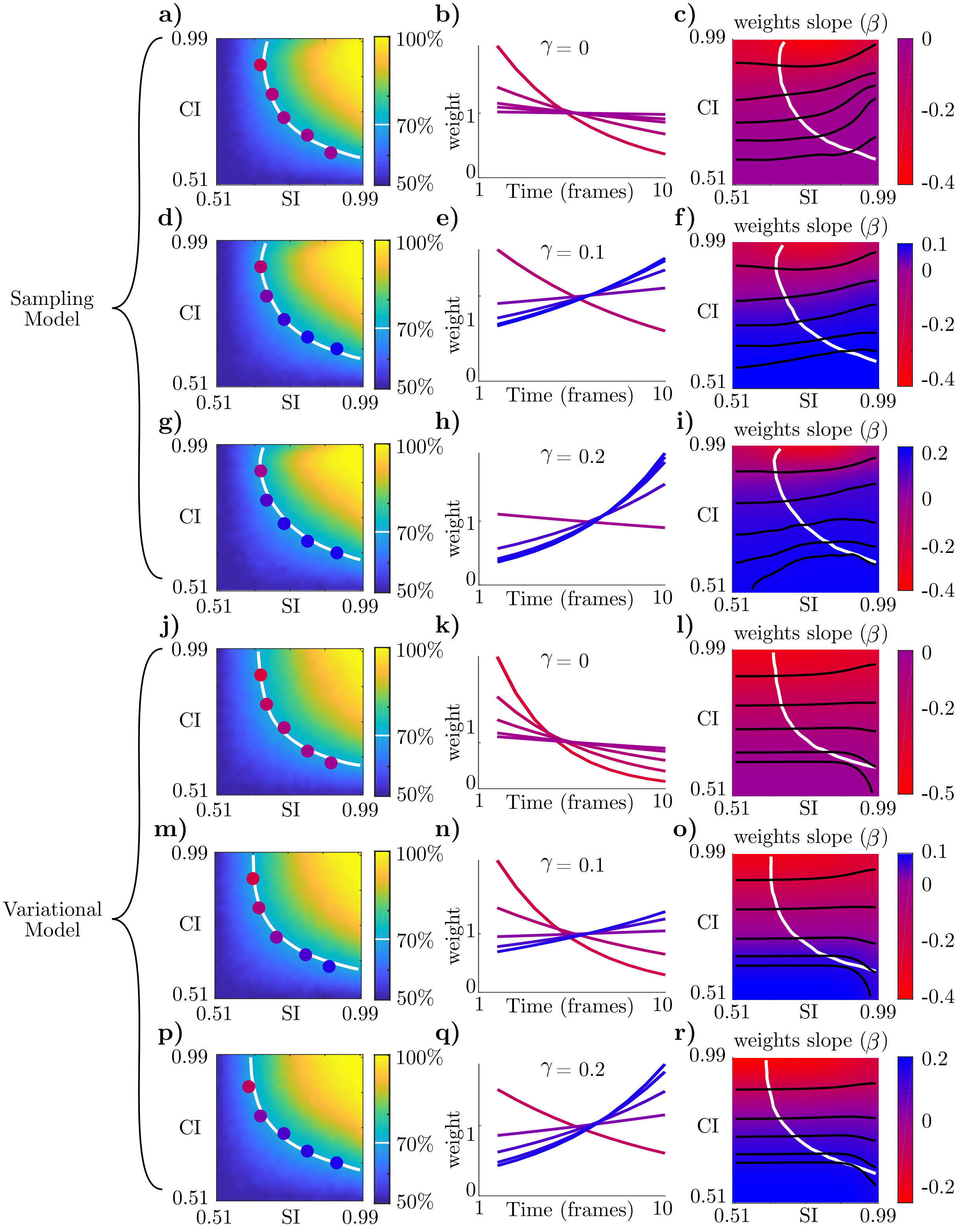


Figure S4: In both models, larger $\gamma$ increases the prevalence of recency effects across the entire task space. Panels are as in Figure 3 in the main text. **a-c** sampling model with $\gamma = 0$. **d-f** sampling model with $\gamma = 0.1$. **g-i** sampling model with $\gamma = 0.2$. **j-l** variational model with $\gamma = 0$. **m-o** variational model with $\gamma = 0.1$. **p-r** variational model with $\gamma = 0.2$.

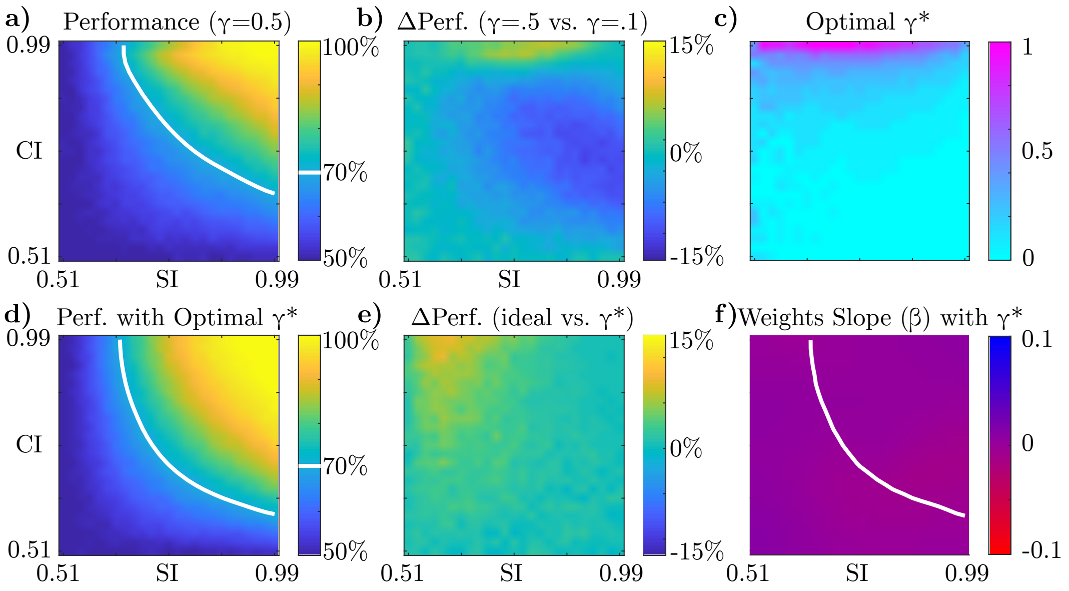


Figure S5: Optimizing performance with respect to $\gamma$ (see also Figure S6). **a)** Sampling model performance across task space with $S = 5$ and $\gamma = 0.5$ (compare with Figure 3c in which $\gamma = 0.1$). **b)** Difference in performance for $\gamma = 0.5$ versus $\gamma = 0.1$. Higher $\gamma$ improves performance in the upper part of the space where the confirmation bias is strongest. **c)** Optimizing for performance, the optimal $\gamma^*$ depends on the task. Where the confirmation bias had been strongest, optimal performance is achieved with a stronger leak term. **d)** Model performance when the optimal $\gamma^*$ from (c) is used in each task. **e)** Comparing the ideal observer to (d), the ideal observer still outperforms the model but only in the upper part of the space. **f)** Temporal weight slopes when using the optimal $\gamma^*$ are flat everywhere. The models reproduce the change in slopes seen in the data only when $\gamma$ is fixed across tasks (compare Figure S4).

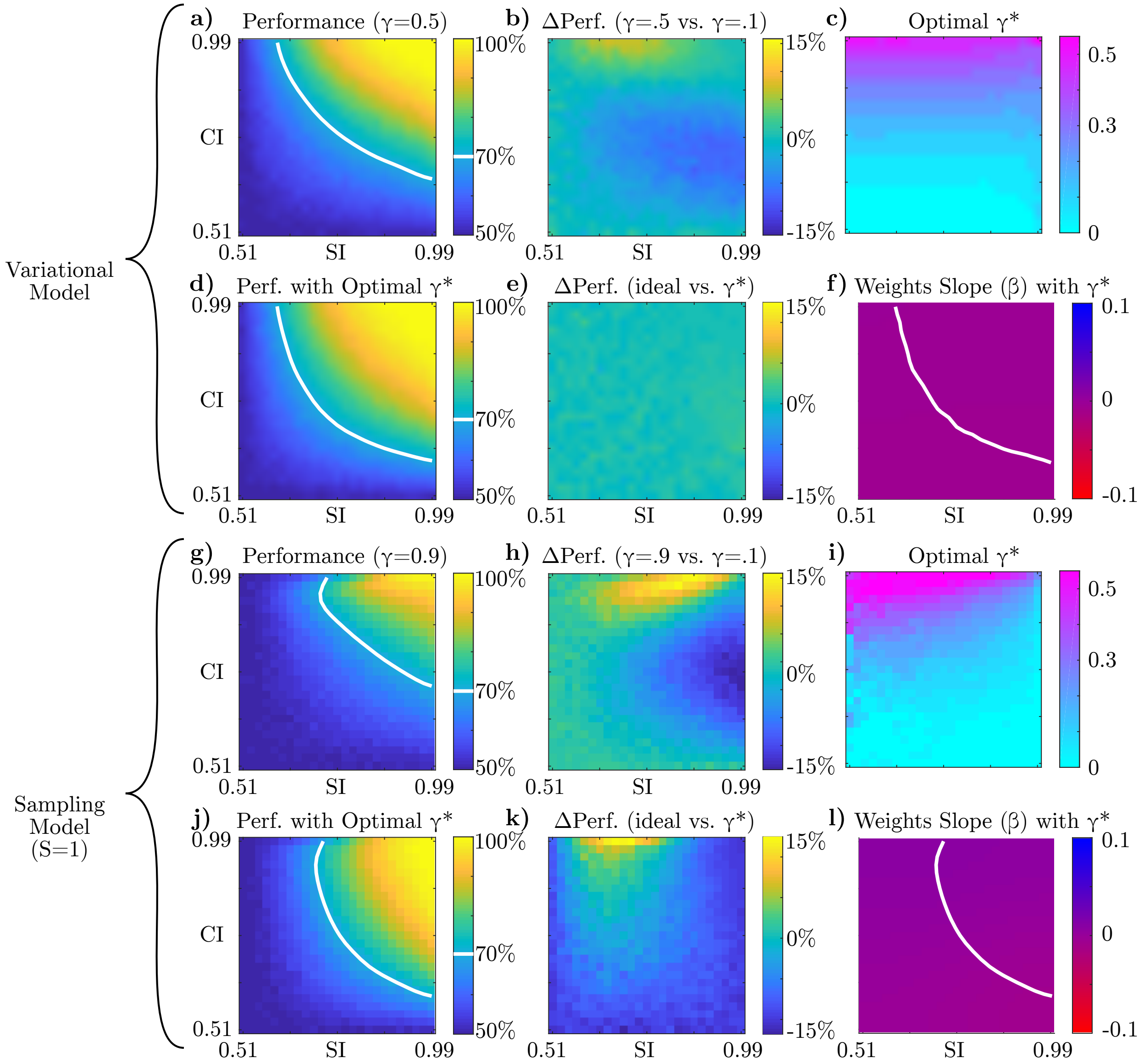


Figure S6: Simulation results for optimal leak ($\gamma$) for two further model variations, panels as in Figure S5. **a-f** Variational model results. As in the sampling model, we see that the optimal value of $\gamma*$ increases with category information, or with the strength of the confirmation bias. **h-l** Sampling model results with $S = 1$ (in the main text and Figure S5 we used $S = 5$). Since the sampling model without a leak term approaches the ideal observer in the limit of $S \to \infty$, the optimal $\gamma^*$ was close to 0 for much of the space in the main text figure. Here, by comparison, $\gamma^* > 0$ is more common because the $S = 1$ model is more biased.

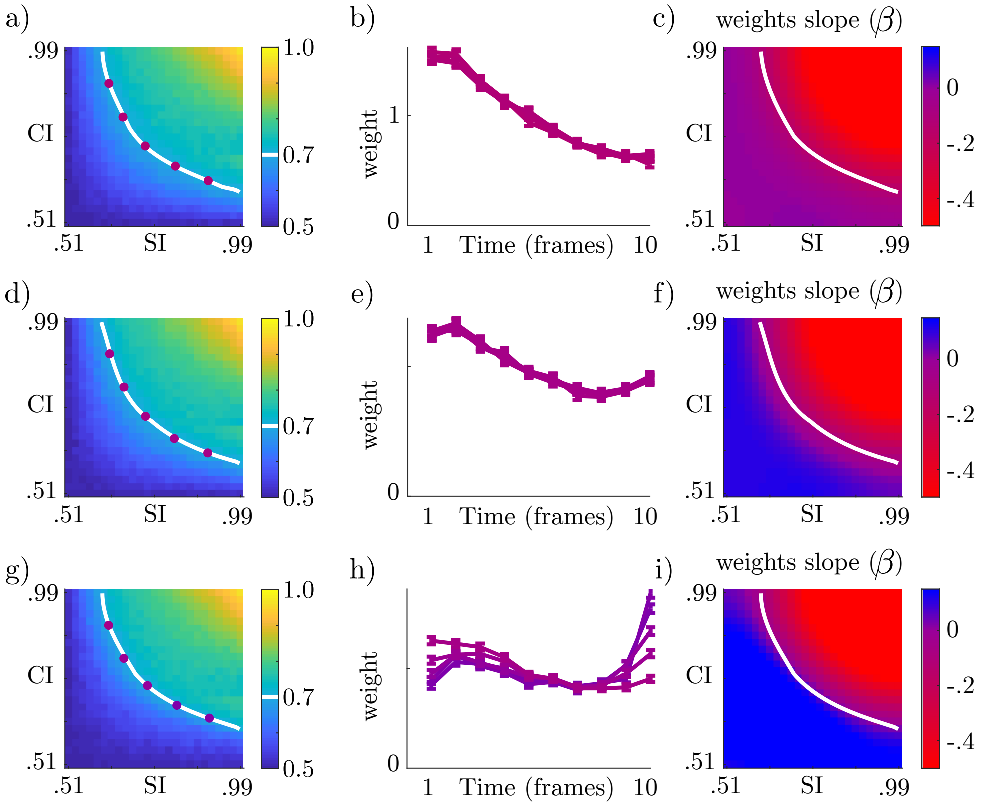


Figure S7: Simulation of bounded integration (ITB) model. **a)** Performance of an ITB model is not differentially modulated by sensory and category information. **b)** ITB consistently produces primacy effects, as in [16]. **c)** The primacy effect becomes more extreme in regions where evidence is stronger, since the bound is hit earlier in the trial. **d-f)** As in (a-c), but with an additional leak term, resulting in less extreme primacy effects and a transition to recency for *difficult* tasks, but no transition from primacy to recency along the iso-performance contour. (Also note the departure from monotonic exponential-like weight profiles). **g-i)** We now vary the leak term, $\alpha$, as a function of category information. This reproduces the qualitative transition from primacy in LSHC to recency in HSLC. As measured by an exponential fit ($\beta$), slopes are matched to those in the confirmation bias models (Figure 3d,g).

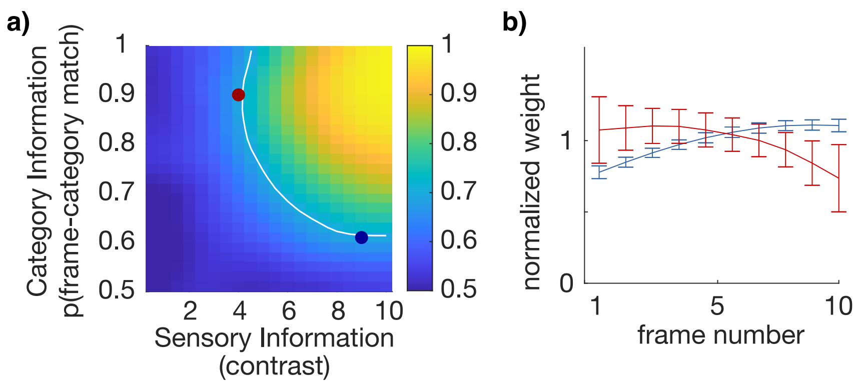


Figure S8: Simulation results on the larger model of [15]. **a)** Performance as a function of sensory information (grating contrast) and category information (probability that each frame matches the trial category). White line is iso-performance contour at 70%, and dots correspond to LSHC and HSLC parameter regimes plotted in (b). Simulation details in the Supplemental Text. **b)** Temporal weights from LSHC and HSLC simulations corresponding to colored points in (a), normalized in each condition so the weights have mean 1. As in the reduced models in the main text, we see a transition from primacy to recency.

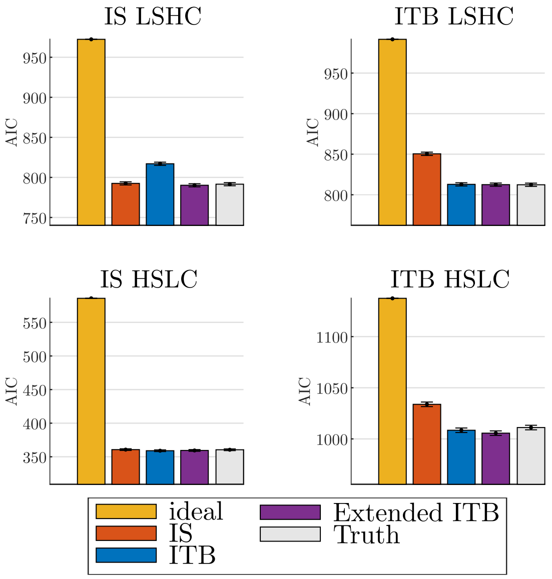


Figure S9: Results of direct model comparison between IS model and ITB model(s) fit to ground-truth data. We employed methods to search the log likelihood landscape of each model despite the stochastic likelihood evaluations of the IS model [24, 2]. Lower AIC indicates better fit. An ideal integrator (gold) and ground-truth (gray) values serve as upper- and lower-bounds, respectively, on plausible AIC values. In all cases, the best fitting model recovered parameters that are as good as the ground truth. The standard ITB model (with positive leak enforced) is distinguishable from the IS model in the LSHC simulation (top row). However, an Extended ITB model that allows for negative leak (purple), fits all data in all conditions as well as the ground-truth. For this reason, we state in the main text that a negative leak is *functionally* indistinguishable from the true IS model. We pursued *parameter comparison* within this Extended ITB model class, rather than *model comparison* between IS and ITB, in the main text.

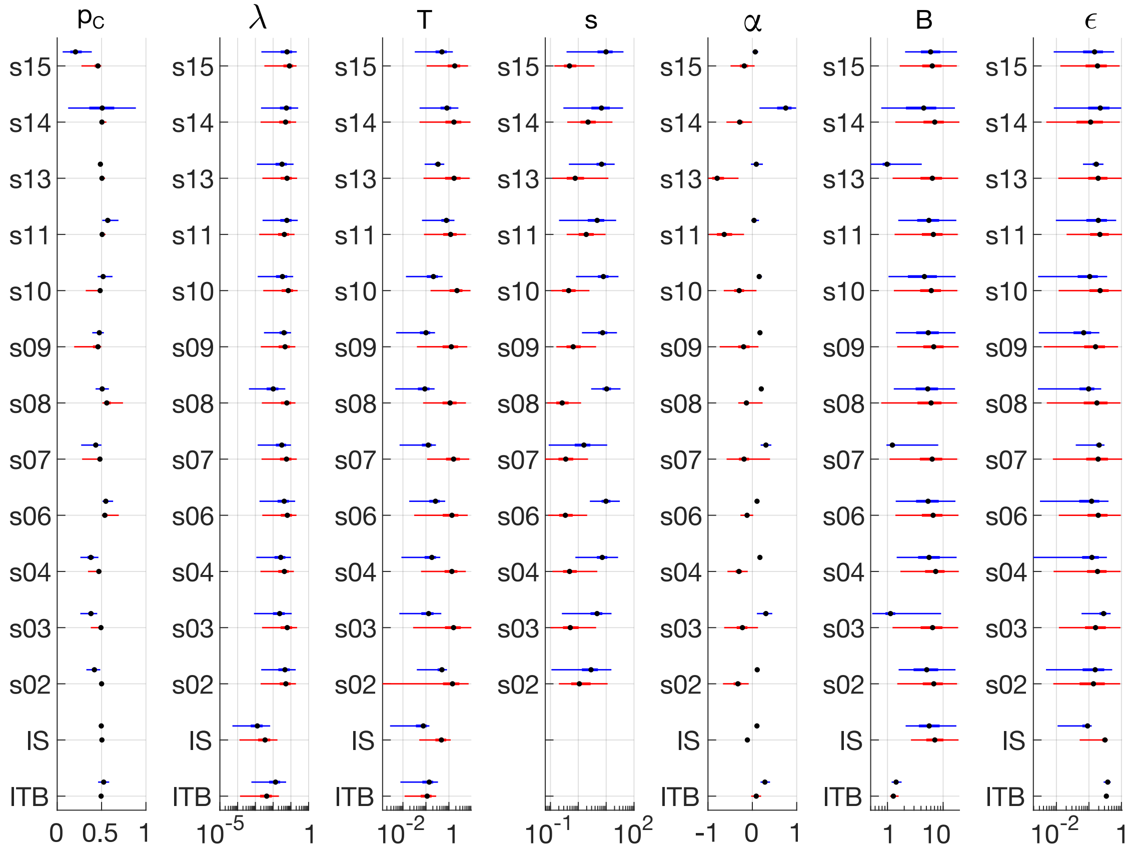


Figure S10: Box and whisker plots of inferred parameter values for each of 12 observers as well as the ground truth models (IS and ITB – see Table S3). Each parameter and observer has two fits, one for the LSHC condition (lower/red) and one for the HSLC condition (upper/blue). Thin lines are 95% posterior interval, thick lines are 50% interval, and points are posterior median. Parameter names are as in the main paper, restated here: $p_C$ = prior over categories, $\lambda$ = symmetric lapse rate, $T$ = decision temperature, $s$ = signal scale (fixed to 1 for ground truth models), $\alpha$ = leak, $B$ = bound, $\epsilon$ = noise.

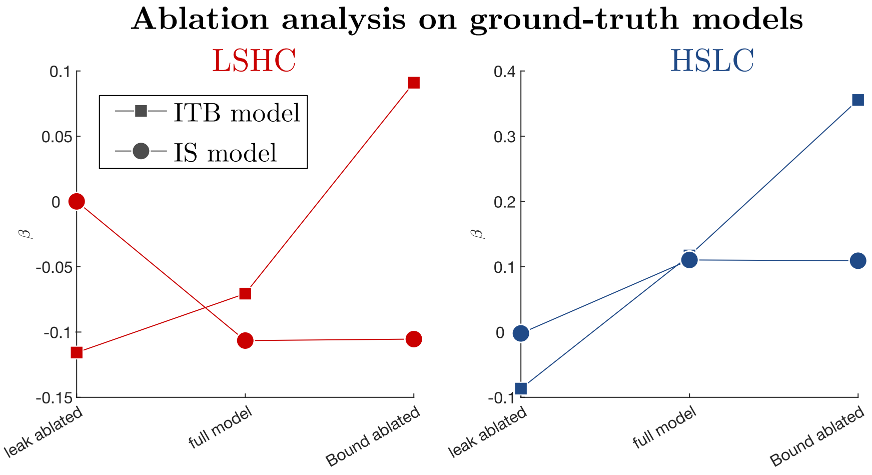


Figure S11: Parameter ablation analysis on ground-truth models. Recall that the ITB model has a primacy effect in the LSHC condition driven by bounded integration. The key signature of bounded integration dynamics is that when the bound is ablated, the leak takes over and it flips to no bias or a recency bias. The key signature of a hierarchical inference model (here, Importance Sampling or IS), on the other hand, is that the primacy bias is unaffected by ablating the bound, but disappears when the leak term is ablated, since a negative leak acts as a confirmation bias. In the HSLC condition (right panel), both models' recency effects are driven by leaky integration. The ITB model's bound competes with the leak, however, so ablating the bound results in exaggerated recency effects, and ablating the leak results in primacy effects. The key signature of a hierarchical inference model, on the other hand, is a recency effect that is unaffected by ablating the bound and that disappears when the leak is ablated.

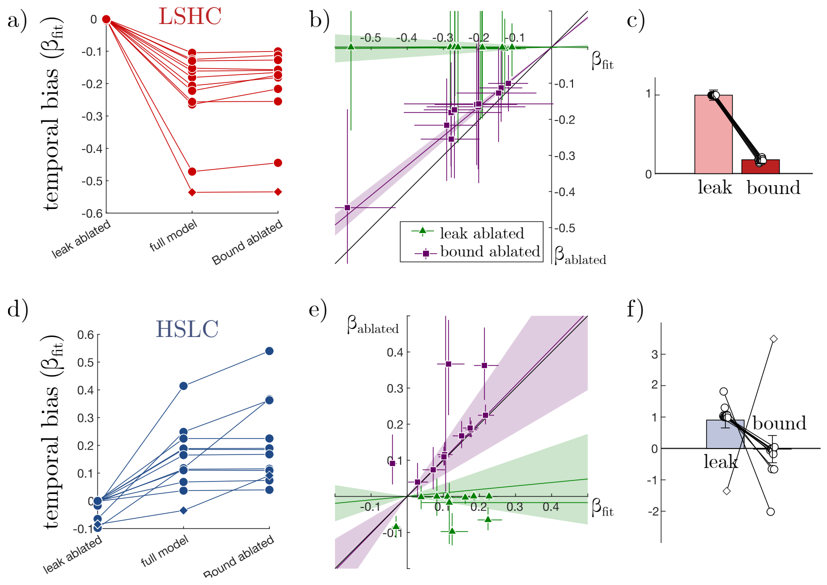


Figure S12: Additional information on fits of the Extended ITB model to empirical data and ablation analyses. **a)** Copy of Figure 6d. Comparing with Figure S11 suggests that primacy effects are largely driven by confirmation-bias dynamics rather than by bounded integration. **b)** Temporal bias of the full Extended ITB model (x-axis) versus the ablated model (y-axis) for each observer and each ablated parameter in the LSHC condition (each observer has two points at the same x coordinate, offset for visualization). We regressed a single slope for each ablated parameter to summarize the fraction of bias in the population explained by the leak parameter (green) or the bound parameter (purple). **c)** Copy of Figure 6e from the main text. The fact that the leak parameter explains 99.4% of the population primacy effects corresponds to the green regression line being nearly horizontal in (b). **d-f)** Same as (a-c) but for the HSLC condition. As in Figure 6, outlier observer in – who had a *primacy* bias in the HSLC condition – is shown as a diamond symbol in panels (a), (d), and (f).

### References

[1] Luigi Acerbi. Variational Bayesian Monte Carlo with Noisy Likelihoods. *arXiv*, 2020.

[2] Luigi Acerbi and Wei Ji Ma. Practical Bayesian optimization for model fitting with Bayesian adaptive direct search. *Advances in Neural Information Processing Systems*, 30:1837–1847, 2017.

[3] William H A Beaudot and Kathy T. Mullen. Orientation discrimination in human vision: Psychophysics and modeling. *Vision Research*, 46:26–46, 2006.

[4] José M. Bernardo and Adrian F. M. Smith. *Bayesian Theory.* Wiley, West Sussex, England, 2 edition, 2000.

[14] David S. Greenberg, Marcel Nonnenmacher, and Jakob H. Macke. Automatic Posterior Transformation for Likelihood-Free Inference. 2019.

[22] Scott A. Sisson, Yanan Fan, and Mark A. Beaumont. Overview of Approximate Bayesian Computation. In Scott A. Sisson, Yanan Fan, and Mark A. Beaumont, editors, *Handbook of Approximate Bayesian Computation*, chapter 1. CRC Press, 2018.

[23] Konstantinos Tsetsos, Juan Gao, James L. McClelland, and Marius Usher. Using time-varying evidence to test models of decision dynamics: Bounded diffusion vs. The leaky competing accumulator model. *Frontiers in Neuroscience*, 6(JUN):1–17, 2012.

[24] Bas van Opheusden, Luigi Acerbi, and Wei Ji Ma. Unbiased and Efficient Log-Likelihood Estimation with Inverse Binomial Sampling. pages 1–89, 2020.

[25] Valentin Wyart, Vincent De Gardelle, Jacqueline Scholl, and Christopher Summerfield. Rhythmic Fluctuations in Evidence Accumulation during Decision Making in the Human Brain. *Neuron*, 76(4):847–858, 2012.
